# Speedy but late: Clock-independent changes in phase in cultivated potato

**DOI:** 10.64898/2026.11.20.767954

**Authors:** Ann Feke, Brieanne Vaillancourt, Kaitlyn Acheson, Joshua C. Wood, Sam Hu, John P. Hamilton, David Hall, C. Robin Buell, Eva M. Farre

## Abstract

One of the mechanisms by which the circadian clock influences plant growth and development is through modulating gene expression timing under light/dark cycles. While wild potato accessions display significant variation in their circadian period, cultivated accessions possess fast clocks independent of their geographic origin. Here, we explore the implications of variation in circadian period on diel gene expression by comparing a short-period cultivar of *Solanum tuberosum* with two wild potato species, the short-period *S. candolleanum* and the long-period *S. chacoense*, through transcriptomics. In all three species, the diel phase of clock-associated genes is well-correlated to the pace of the clock. However, *S. tuberosum* displays a significant genome-wide delay in gene expression phase despite its short circadian period. This delayed phase was also observed in both diel leaf movements and sucrose content. Reconstruction of the transcriptional network indicates that DOF and REVEILLE transcription factors play a major role in diel expression patterns in these species. In modern cultivated potatoes, truncated alleles of the DOF transcription factor CYCLING DOF FACTOR 1 enable early tuberization. We find that CDF1 is able to repress *RVE1a* expression *in vivo.* We propose that truncated CDF1, in combination with its downstream targets, leads to delays in gene expression that are decoupled of circadian control.

## Introduction

Due to their sessile nature, plants must be able to adapt to the wide array of environmental conditions they are exposed to throughout the year. Depending on the geographic range of a given plant species, individuals may be exposed to varying ranges of daylength, precipitation and temperature which often correlate strongly with the seasons. The circadian clock has been found to affect a significant proportion of the physiological and developmental processes performed by the plant (Greenham and McClung, 2015), and is hypothesized to help plants anticipate and adapt to seasonal and latitudinal daylength changes.

Several studies have investigated the prevalence of variation in circadian clock traits in both natural populations and domesticated cultivars; see (Oravec and Greenham, 2022; González-Delgado et al., 2025; Feke and Farré, 2026) for recent reviews on this topic. This variation has largely been divided into two major categories: variation in circadian period that correlates well either with latitude or elevation in natural populations, or targeted changes in clock-associated genes to improve resilience, yield, or other beneficial traits in domesticated species. The circadian period is measured under constant environmental conditions and reflects the genetically determined speed of the endogenous circadian oscillator. Since most plants are never exposed to constant light, it raises the question of the role of the circadian period under natural light/dark cycles. Under diel cycles, the resetting of the clock at dawn and dusk leads to consistent 24-hour periods. Natural variation of the endogenous free running period has been associated with changes in phase under light/dark conditions of some circadian regulated traits, such as gene expression, in a few plant species (Müller et al., 2016; Muller et al., 2018; Muranaka et al., 2022). However, we still lack a comprehensive understanding of the interaction between variation of the circadian period and physiological oscillations in diel conditions.

In potato, while domesticated species exhibit consistently short circadian periods, wild potato species vary significantly in the speed of their clocks (Hardigan et al., 2017). The short circadian period of domesticated potatoes, even in accessions grown at high latitudes, differs with what has been observed in closely related tomato, where a long circadian period has been associated with domestication (Müller et al., 2016). The slow clock of cultivated tomatoes leads to a late phase of gene expression and has been linked to higher chlorophyll content under the longer daylengths that occur at higher latitudes. Therefore, we investigated the significance of the short circadian period in cultivated potatoes and the role of circadian period variation in wild tuber bearing *Solanaceae*.

We compare the genome-wide diel expression patterns of the domesticated short-period cultivar *S. tuberosum* cv. Atlantic with two wild species, S*. candolleanum*, which also possesses a short circadian period, and *S. chacoense*, which has a significantly longer circadian period. We performed this analysis under two photoperiods, a tuber-inductive short-day photoperiod and a tuber-inhibitory long-day photoperiod. We hypothesized that shorter circadian periods lead to early phasing of gene expression and physiological responses under diel conditions. Our results indicate that while the timing of expression of genes associated directly with clock function are well-correlated with the circadian period in all three species, genome-wide phasing is only correlated with circadian period in wild species. In contrast, despite their short circadian periods, the majority of genes in the domesticated potato exhibit significant phase delays compared to both wild species, suggesting an alternate mechanism of circadian adaptation outside of modulating the free running period in potato cultivars. Through construction of a gene expression network, we propose a cascading mechanism that causes a lag in rhythmic expression in cultivated potatoes in spite of their short circadian period.

## Results

### Selection of tuber-bearing *Solanum* species

To investigate the impact of circadian clock variation in potato we selected three species from the tuber bearing *Petota* clade, two wild species with naturally-derived differences in circadian periods, and a cultivated *S. tuberosum* cultivar that has been developed for northern hemisphere climates significantly different from its ancestral range. Both the wild *S. candolleanum* accession and *S. tuberosum* have short circadian periods of around 20 hours, while the diploid inbred clone of the wild *S. chacoense* has a long circadian period of around 25 hours (Fig. 1A). *S. candolleanum* has been hypothesized as the progenitor species of the domestic *S. tuberosum* (Spooner et al., 2005). Thus, these accessions enabled us to investigate both the effects of natural variation in circadian period and the effects of domestication on rhythmic gene expression.

**Figure 1.**
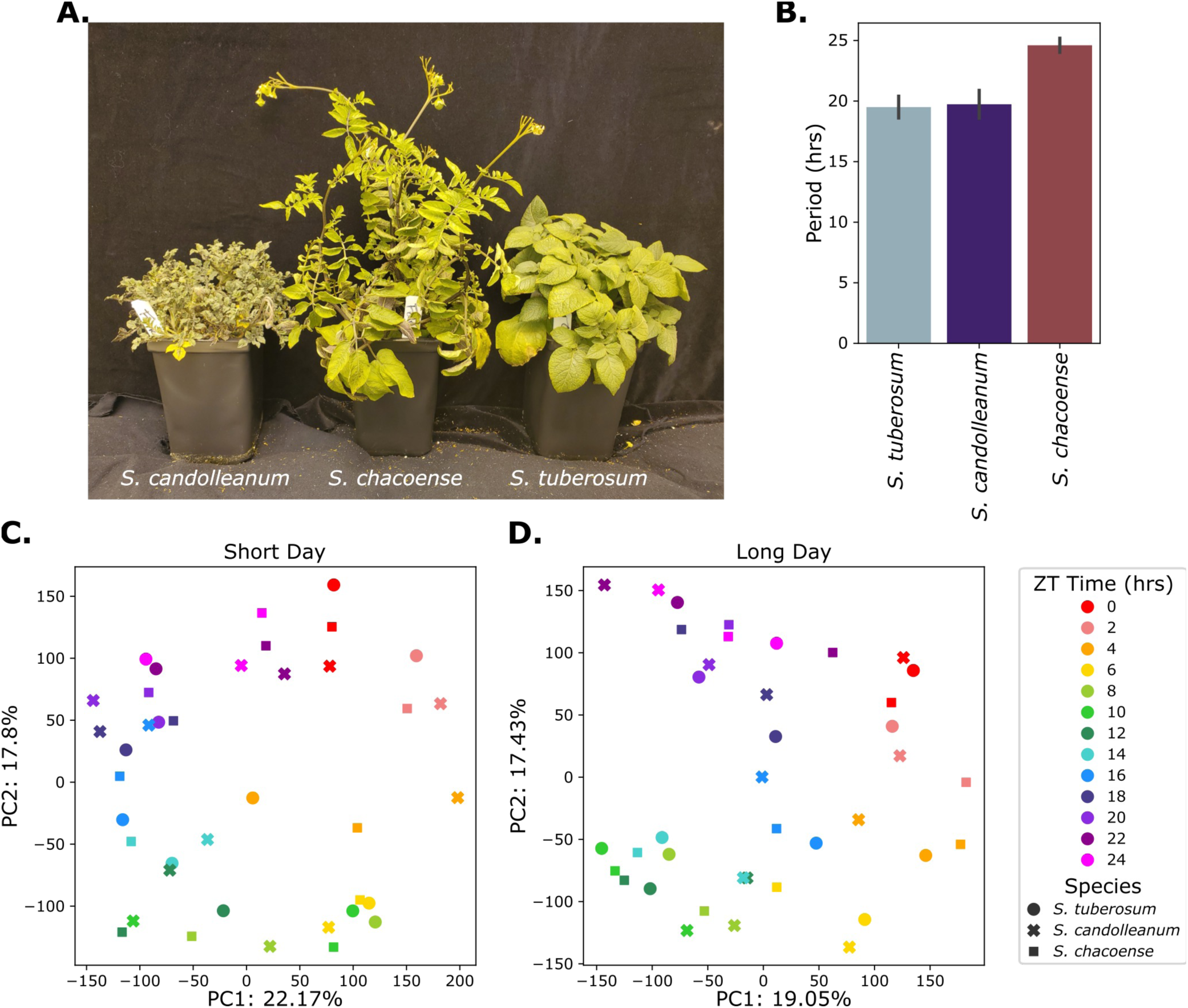
Differences in expression between three tuber-bearing *Solanum* species is driven by time, rather than species. A. Photographs of seven-week-old *S. candolleanum* (left), *S. chacoense* (center) and *S. tuberosum* (right) plants grown under long day conditions. B. Circadian period as measured by delayed fluorescence for *S. tuberosum*, *S. candolleanum* and *S. chacoense.* C-D. Principal component analysis of shared expressed syntelogs under C. short day (12L/12D) and D. long day (16L/8D) conditions in *S. tuberosum* (circles), *S. candolleanum* (crosses) and *S. chacoense* (squares). Time is represented by color, with warmer colors representing those close to dawn, and cooler colors representing those closer to dusk.

### Genomic resources for S. candolleanum and S. chacoense

To compare across our accessions, we generated new or improved genomic resources for our wild *Solanum* accessions. For both *S. chacoense* and *S. candolleanum,* we utilized Oxford Nanopore long reads along with scaffolding using Hi-C datasets, synteny with DM v6.1 (Pham et al., 2020), and concordance with genetic maps to generate chromosome-scale genome assemblies. For *S. chacoense,* we generated a 676.2 Mb assembly with an N50 scaffold length of 53.1 Mb; for *S. candolleanum,* we generated a 699.8 Mb assembly with an N50 scaffold length of 55.3 Mb (Table S1). Benchmarking Single Copy Orthologs (BUSCO) analysis with both the genomes and predicted proteomes (Table S2, Table S3) coupled with the high contiguity metrics of the assemblies are consistent with high-quality chromosome-scale genome sequences. These new resources, along with the haplotype resolved resources for *S. tuberosum* cv. Atlantic (Hoopes et al., 2022) facilitated mining our transcriptomics datasets on their cognate genomes.

### Identification of rhythmic genes

To investigate the effects of variation in circadian period on the phase of gene expression under diel conditions, we performed a transcriptomic profiling experiment on each of the three potato species under both short (12-hour light/12-hour dark) and long day (16-hour light/8-hour dark) conditions. We used synteny to determine which genes had high-confidence syntenic orthologs across all three species (Fig. S1A, Datasets S1-7). To query the extent of time-dependent gene expression under diel conditions, we first performed principal component analysis (PCA) of our transcriptome data. We found that time, not species, drove the majority of separation between our samples in both photoperiods, suggesting significant conservation of time-dependency of gene expression in our dataset (Fig. 1B-C). We then quantified rhythmicity using the JTK algorithm (Hughes et al., 2010)(Datasets S8-13). We found that under short days 2,398, or 16% of expressed syntelogs, were rhythmic in all three species (Fig. S2A). A slightly smaller percentage of genes were rhythmic under long days as compared to short days in *S. tuberosum* and *S. candolleanum*, while *S. chacoense* had a larger percentage of rhythmic genes under long day conditions (Fig. S2B-D). Genes displaying rhythmic expression in all three species were enriched in functions related to circadian rhythms and well-characterized clock outputs such as light responses, photosynthesis and primary metabolism (Table S4). Taken together, this suggests we selected a functionally relevant set of genes for our species comparisons.

### The phase of circadian clock associated genes correlates with free-running circadian period

One of the predicted impacts of variation in circadian period is a change in phasing of rhythmic gene expression under diel conditions, with shorter circadian periods leading to earlier phases and vice versa (Aschoff and Pohl, 1978). To investigate the impact of natural variation of the circadian period on phase, we first investigated the expression of a set of 26 conserved circadian clock-related genes that were expressed and rhythmic in all three species (Table S5). Overall, the patterns of diel expression for these genes were highly similar across the three genotypes (Figs. 2, S3-5). We observed a correlation between the phase and the circadian period such that the short period *S. tuberosum* and *S. candolleanum* had, on average, significantly earlier phases (0.9 and 0.76 hours respectively) than *S. chacoense* (Wilcoxon signed-rank test p-value < 0.05) (Figs. 2A, S4A). We observe a similar tendency under long day conditions, such that *S. chacoense* was phased later than both *S. tuberosum* and *S. candolleanum* (0.5 and 0.9 hours later on average, respectively, Wilcoxon rank-sum test p<0.01) (Figs. S3A, S5).

**Figure 2.**
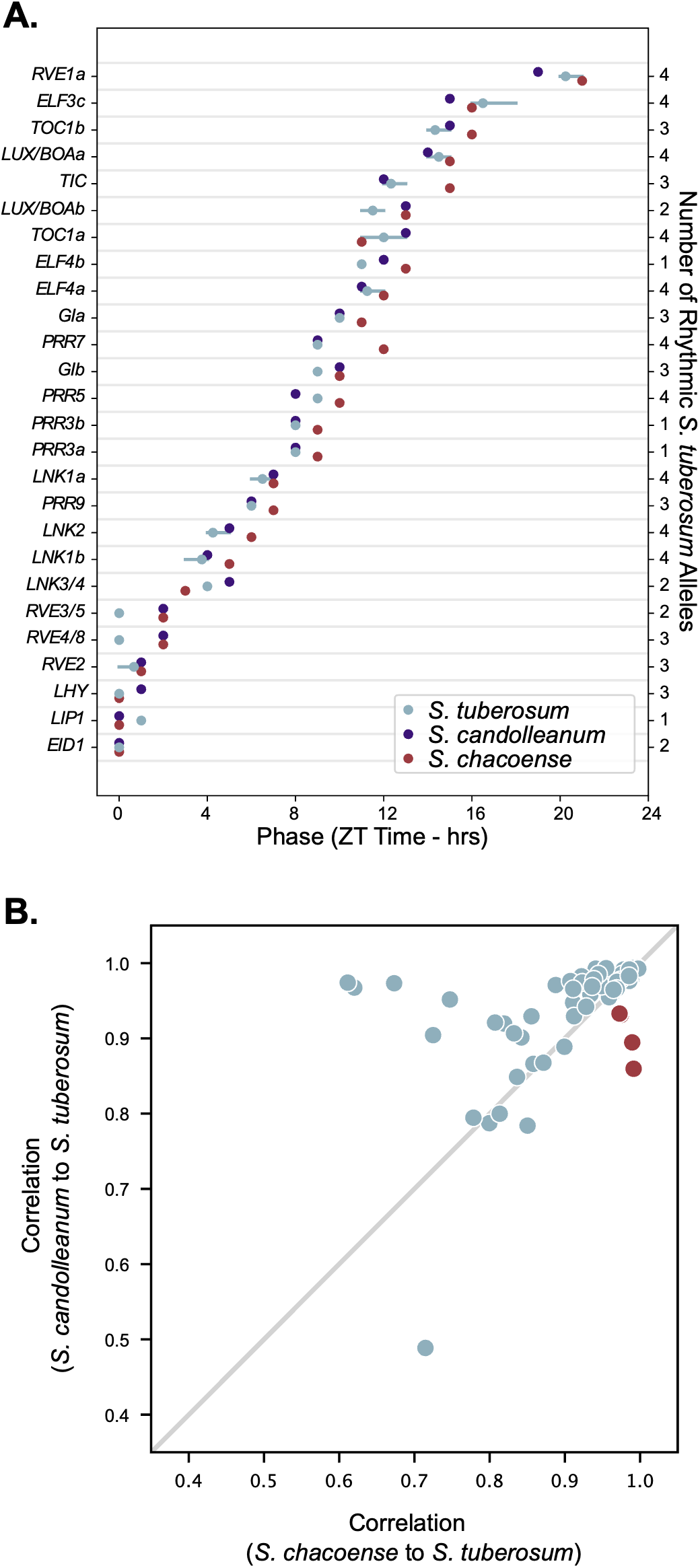
Circadian period correlates with clock gene phase under short day conditions. A. The phase of expression of rhythmic clock genes in *S. tuberosum* (blue), *S. candolleanum* (purple), and *S. chacoense* (red) under short day conditions. As the *S. tuberosum* assembly was haplotype-resolved, anywhere from 1-4 alleles can be included for each *S. tuberosum* datapoint. For each species, the circles represent the mean phase of all expressed and rhythmic alleles, while the horizontal lines represent the range of phases for each gene. The number of alleles that is included for each gene is listed on the right y-axis. B. Pearson correlations between the gene expression patterns of rhythmic clock genes *S. tuberosum* compared to either wild species under short day conditions. *RVE1a* is highlighted in red.

In accordance with the similarities in phase, the expression patterns of the majority of *S. tuberosum* clock associated genes displayed significantly higher correlation with those from *S. candolleanum* than those of *S. chacoense* (Figs. 2B, 3B). One of the few exceptions is *REVEILLE 1a* (*RVE1a*), whose rhythmic expression pattern was more similar to that of *S. chacoense* than *S. candolleanum* under both photoperiods. These observations support a role of the circadian clock in determining the phase of expression under light dark cycles.

**Figure 3.**
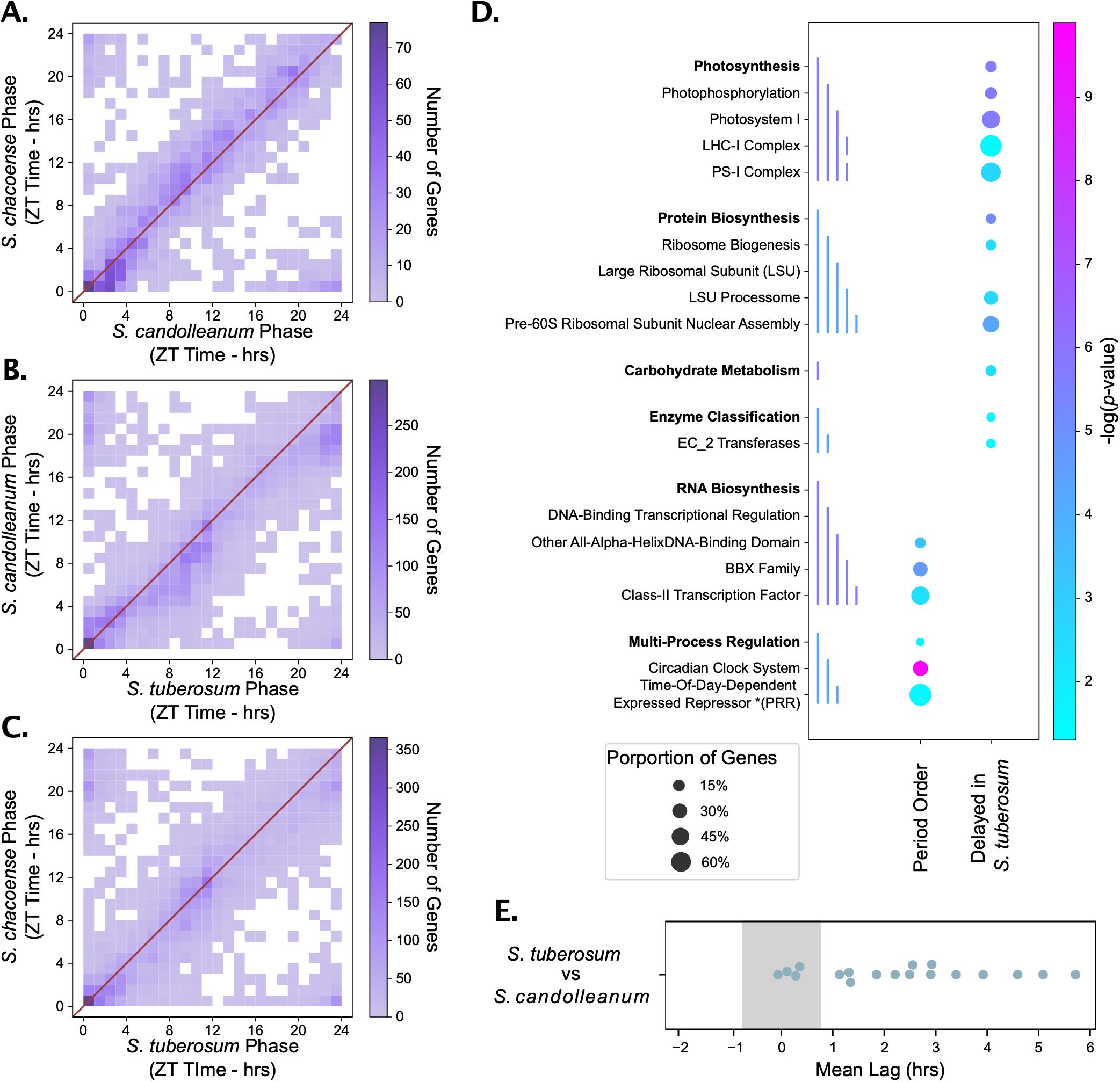
The phase of rhythmic gene expression genome-wide correlates with circadian period in wild species but not domestic species. A-C. Two-way histograms of phase across all genes that were rhythmic under short day conditions in both A. *S. candolleanum* and *S. chacoense* B. *S. candolleanum* and *S. tuberosum*, or C. *S. chacoense* and *S. tuberosum.* The red lines represent the points where phase is equal in both species. *S. chacoense* is significantly delayed in comparison to *S. candolleanum*, while *S. tuberosum* is significantly delayed in comparison to both wild species (Wilcoxon Signed Rank Test *p*-value<0.001). D. Mapman enrichment for genes that have phases that correlate with their circadian period in all three species ("Period Order") and those which have a delayed phase in *S. tuberosum* ("TUB Late"). The size of the circle represents the percentage of the number of rhythmic genes with that mapman term that were found in the phase category, and color represents the –log(p-value) of the enrichment. Colored lines on the left side of the graph represent the hierarchical relationship between related mapman terms. E. Average lag of all the genes within *S. candolleanum* and *S. tuberosum* modules. Positive values are those where *S. tuberosum* is delayed compared to *S. candolleanum*.

### Circadian period and genome-wide gene expression phase only correlate in wild species

To determine whether the natural variation in circadian period between the two different wild *Solanum* species followed the trend observed in the circadian associated genes, we compared the phases of syntelogs rhythmic in both *S. candolleanum* and *S. chacoense* (Fig. 3A). Under short days, the average phase was approximately 0.4 hours earlier in S*. candolleanum* than in *S. chacoense* and ∼30% of rhythmic genes displayed a delay in *S. chacoense* of at least 2 hours. Although both species exhibited significantly delayed phasing in long days as compared to short, no statistically significant difference in phase between these two genotypes was measured under long days (Fig. S6). In contrast, we observed significant delays in the average gene expression between *S. tuberosum* and *S. candolleanum* of 0.9 and 1.2 hours in short and long days respectively (Figs. 3B, S6C). In short days, about 43% of genes had a delayed phase of at least two hours in *S. tuberosum*. We also observed a significant delayed phase in *S. tuberosum* with respect to *S. chacoense* (∼0.7 hours), but only under the short photoperiod (Figs 3C, S6D). In this condition, about 34% of rhythmic genes had at least a two-hour delay in expression in *S. tuberosum*. These results suggest that, while the domestic cultivar *S. tuberosum* has a fast circadian clock, a period-independent mechanism is being employed to delay the phases of gene expression under diel conditions. Genes with a late phase in *S. tuberosum* were enriched in photosynthesis and ribosome biogenesis genes indicating that this late phase might have an impact on plant growth (Fig. 3D). In contrast, genes with phase correlated to the endogenous circadian period were enriched in circadian clock associated genes and BBX transcription factors, which include the *COL* genes, indicating that these genes may be directly regulated by core clock components (Fig 3D).

### Co-expression modules identify functional targets of domestication-induced phase delays

To investigate the potential mechanism and implications of the delay in phase in *S. tuberosum*, we generated co-expression modules based on the expression patterns of genes in pairs of species. We had previously determined that time was a major driving factor in the first two principal components in our PCA (Fig. 1B-C). We therefore generated our co-expression modules with genes that had a significant impact on either the first or second principal component. This strategy enabled us to include any gene with time dependent expression patterns without restricting to those with the sinusoidal pattern required for rhythmicity according to the JTK algorithm. We focused on comparing *S. tuberosum* with *S. candolleanum* under short days to identify genes that are delayed in *S. tuberosum* in a clock-independent manner. Using this methodology, we generated 18 co-expression modules for this pair of species, encompassing 6,667 syntelogs, or ∼44% of all expressed syntelogs (Table S6). The average gene expression pattern for each of these modules had an apparent rhythmic pattern, although several modules appeared to have more than one peak throughout the day (Fig. S7, left column). These results indicate that our co-expression modules are likely representative of major rhythmic gene expression patterns.

We then quantified the phase difference for each of these modules by calculating the lag correlation between species for each gene within a module. We determined that a module had a difference in phasing between species if the magnitude of the average lag correlation was greater than one hour. Using this methodology, we identified that 14 of the 18 modules were delayed in *S. tuberosum*, while the remainder exhibited no differences in phase (Fig. 3E). Most (11 out of 14) of these delayed modules had elevated gene expression during the second half of the day (∼ ZT8) in *S. tuberosum* in comparison to *S. candolleanum* (Fig. S7, left column). These results are consistent with our prior JTK results and show that the delay in phase in *S. tuberosum* versus its wild ancestor was robust to changes in methodology.

We observed that genes in modules were also co-expressed under long days (Fig. S7, middle column) supporting common regulatory mechanisms in both photoperiods. Under long days, 14 of the 18 modules displayed apparent biphasic expression patterns, 12 of which were among the modules that exhibited phase delays in *S. tuberosum* under short days. These biphasic rhythms also appeared to have a higher amplitude in *S. tuberosum* than in *S. candolleanum* in several modules (modules 5, 8, 9, 13, 15, 16 and 17) (Fig. S7, middle column), which could explain why fewer genes were detected as rhythmic under long days than under short days since rhythmicity detection using JTK relies on the pattern fitting a sinusoidal model (Fig S2B-D) (Hughes et al., 2010). In contrast to short days, under the long photoperiod, most modules had no strong differences in phase between both species. The exception was module 12, in which the *S. tuberosum* genes had a maximum of expression during the light period and the *S. candolleanum* genes during the night under long days, but no difference in expression under short days.

To understand the potential physiological implications of our co-expression network we performed functional enrichment analyses (Fig. S7, right column). Genes in modules (1, 2 and 3), which did not display significant shifts in expression between *S. tuberosum* and *S. candolleanum*, included genes involved in light signaling and circadian control and confirm the correlation between diel phase and the endogenous circadian period identified during the JTK analyses (Fig 2A; Fig. S7, right column). For example, module 2, with an early morning peak of expression, was enriched in genes related to light perception and signaling, such as the phytochrome and UV-B signaling pathways. Module 2 also included morning expressed circadian associated genes, while modules 1 and 3 included genes peaking in the afternoon and evening. These unshifted modules also contained several of the BBX transcription factors, including several *CONSTANS-LIKE* genes. These results further support the important role of the clock in mediating the phase of expression of light signaling components and the conservation of clock function between cultivated potato and its wild ancestor.

Genes involved in the light reactions of photosynthesis, including photosystem assembly were enriched in modules 9-11, all of which exhibited a phase delay in *S. tuberosum* (Fig. S7). These genes displayed extended elevated expression during the second half of the light period under short days (Fig. S8). Interestingly, module 9 also contained more than half of all nuclear encoded genes involved in plastid ribosome biogenesis, suggesting coordinated translation of photosystem-associated proteins. These results indicate that *S. tuberosum* and *S. candolleanum* may display differences in the capacity of the photosystem to repair their photosystems throughout the day.

Several shifted modules displaying elevated late-afternoon expression were enriched for other genes involved in growth-related processes. Modules 8 and 13 were enriched in cell wall organization related genes and include ∼25% of all the genes annotated as being involved in hemicellulose biosynthesis or modification (Fig. S7) as well as four cellulose synthase genes. Module 13 was also enriched in genes involved in the microtubular cytoskeleton and cellulose synthase complex trafficking. Module 11 was enriched in genes involving DNA replication and includes half of all the Minichromosome Maintenance (MCM) DNA helicase genes found in our syntelogs. Finally, Modules 14 and 16, which had similar expression patterns, were enriched in genes encoding for histones (Fig. S7). The similarity of expression of these modules (8, 13, 11, 14 and 16) supports the tight regulation of cell wall and cell cycle related processes during plant growth (Soni and Bacete, 2023).

Under long days, Module 12 displayed the largest differences between the phase of *S. tuberosum* and *S. candolleanum*, although it showed similar expression patterns under short-day conditions (Fig. S7). Ribosome biogenesis genes were strongly enriched in this module. In addition, this module contained multiple genes hypothesized to be involved in translation initiation such as the EF1B eEF1A-GDP-recycling complex and the pre-initiation complex (PIC). We had previously observed that ribosome biogenesis genes were among the genes with strongest phase difference between short and long photoperiods in *S. tuberosum* (Feke et al., 2025). The genes in module 12 also displayed strong differences in phase between photoperiods in *S. chacoense* but not in *S. candolleanum*, leading to similar expression patterns of protein biosynthesis genes under long days in *S. tuberosum* and *S. chacoense* (Fig. S9).

### Potato species display photoperiod-dependent differences in biomass, leaf starch and the timing of leaf movement

The circadian clock regulates a wide range of physiological responses in plants. Natural variation in the circadian period has been proposed to optimize growth and reproduction under different photoperiods (Müller et al., 2016; Greenham et al., 2017; Rubin et al., 2017; Feke and Farré, 2026). Since our expression analyses also indicated that cell growth and light reactions of photosynthesis might display differences in timing we quantified several physiological responses in our three genotypes.

We measured photosynthetic parameters using chlorophyll fluorescence under two light intensities and two photoperiods (Figs. S10-11). Under our block light regime, photosystem II efficiency decreased during the second half of the light period, particularly under long-day conditions. Accordingly, non-photochemical quenching (NPQ(T)) (Tietz et al., 2017) increased during the day (Fig. S10). This increase was stronger under high light, indicating it is likely related to accumulation of photodamage. However, we did not observe consistent differences in the timing of these changes between our genotypes (Fig. S11).

It has been proposed that modulation of the circadian period could influence growth under different daylengths and we observed cell division genes displaying differences in phase between our genotypes (Fig. S7). Therefore, we first analyzed biomass accumulation in our three potato species under both long and short photoperiods (Fig. S12). We only observed significant differences in the biomass of the whole plant in the domesticated *S. tuberosum*, wherein long-day growth leads to a reduction in biomass, particularly in the aerial tissues. In contrast, both wild species exhibited no significant changes in the biomass of the entire plant or the aerial tissue. *S. candolleanum* did exhibit a reduction in the below-ground tissue under long days despite not tuberizing in either condition during the short duration of our biomass experiment.

Leaf movement is a circadian controlled trait caused by changes in timing of cell expansion in the leaf petioles (Dornbusch et al., 2014). We measured leaf movement during the light period in the same experiments as the photosynthetic measurements (Fig. 4). Under light/dark cycles, potato leaves open during the day but start closing at the end of the light period by moving upwards. Under short days in both light intensities, the leaves begin to close earlier in *S. candolleanum*, followed by *S. tuberosum* and then *S. chacoense*. This difference in timing between *S. tuberosum* and *S. candolleanum* is consistent with the genome-wide delays in phasing that we observed in our transcriptomic studies. However, under long days we only observed a difference in phase between *S. tuberosum* and *S. candolleanum* under the low-light condition. It is possible that the combination of a long photoperiod and high light promotes earlier upwards leaf movement due to increased light stress.

**Figure 4.**
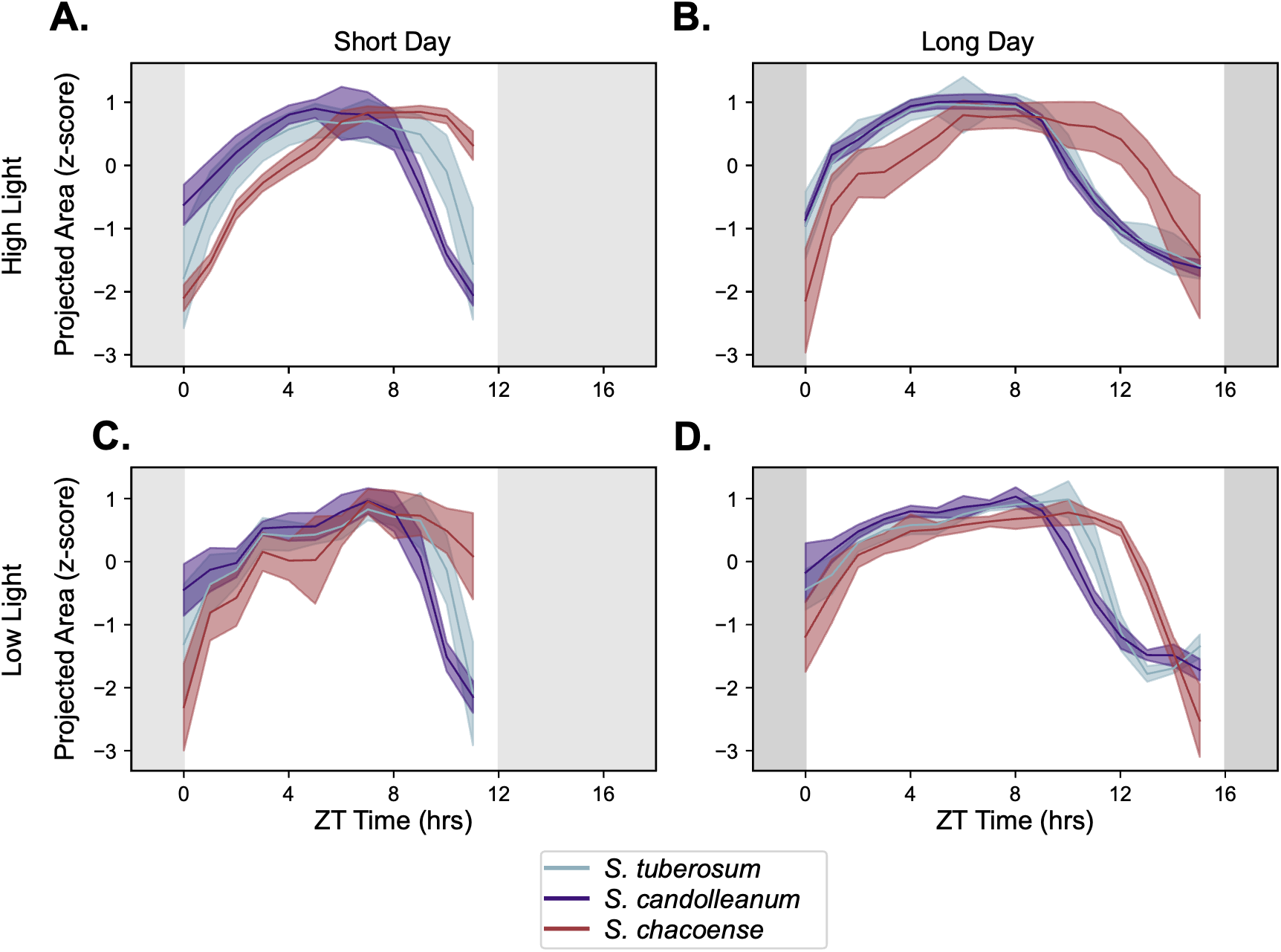
*S. tuberosum* has a delayed phase of leaf movement. Plots of leaf position in A-B. 1000 mmol m^-2^ s^-1^ and C-D.180 mmol m^-2^ s^-1^ under A, C) short days and B, D) long days for *S. tuberosum* (blue), *S. candolleanum* (purple) and *S. chacoense* (red). Lines represent the mean of 16 independent samples while shaded regions represent the standard deviation. Grey boxes represent nighttime conditions.

Rhythmic changes in leaf transitory starch metabolism have also been associated with optimized plant growth (Sulpice et al., 2009). The clock affects the timing of leaf transitory starch degradation at night, and some *A. thaliana* (Arabidopsis) short period mutants degrade starch more quickly during the dark period (Graf et al., 2010). Although we saw no clear differences in phase, we observed differences in the amplitude of starch oscillations between species and photoperiods (Fig. S13). Under short days, all three species displayed clear oscillations, however, *S. candolleanum* had higher leaf starch content at dusk than the other two species. Under long days, *S. tuberosum* maintained similar starch levels as in short days. In contrast, under long days the maximum starch content was reduced in *S. candolleanum*, while *S. chacoense* starch levels failed to oscillate and remained elevated throughout the 24-hour time course. Leaf sucrose content oscillated in all three accessions under long days but only in *S. tuberosum* and *S. candolleanum* under short days. The sucrose content maximum was delayed in *S. tuberosum* in comparison to *S. candolleanum* under both photoperiods (Fig. S13). Our shifted modules contained several carbohydrate-related genes (Table S7). Three sucrose synthase and two neutral invertase encoding genes displayed significant lagged expression in *S. tuberosum* indicating that there could be shifts in the timing of leaf metabolism in domesticated potato (Figs. S14-15).

While collecting our tissue for transcriptomic analysis, we observed that all three species were able to tuberize under the short photoperiod, although only *S. tuberosum* produced tubers under the long photoperiod of 16h light/8h dark. To evaluate the photoperiod sensitivity of tuberization, we grew plants under slightly shorter days of 14h light/10h dark. Under these conditions *S. tuberosum* and *S. chacoense* produced tubers but *S. candolleanum* did not (Fig 5A). These results indicate that our two wild species display differences in the critical photoperiod for tuberization induction. While the presence of an unregulated CDF1 protein in cultivated potatoes leads to tuberization under longer photoperiods (Kloosterman et al., 2013), it is still unclear what modulates tuberization time variation in wild potatoes.

**Figure 5.**
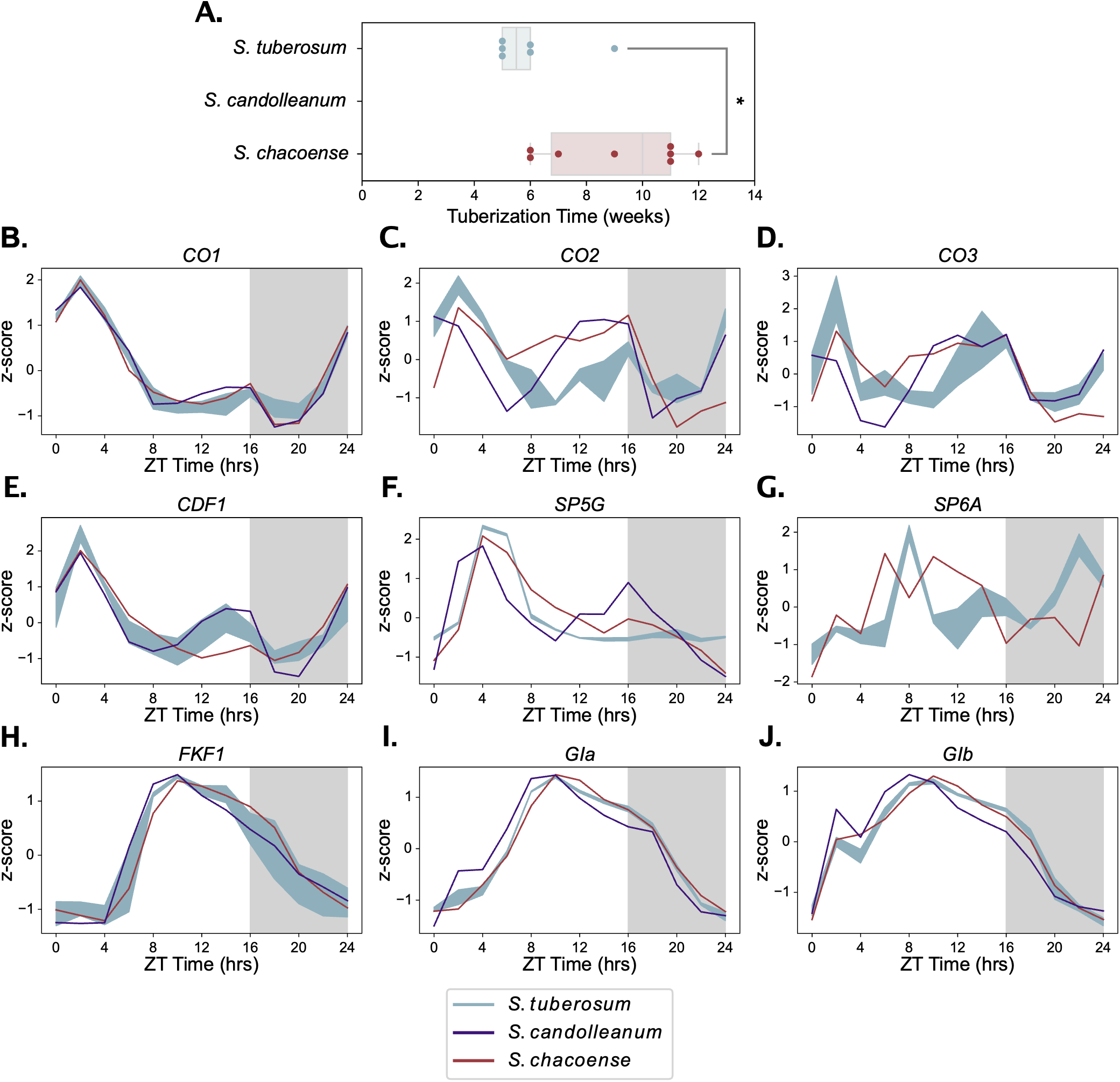
Differences in photoperiodic regulation in three tuber-bearing *Solanum* species. A. Tuberization time measured in weeks after planting for *S. tuberosum, S. candolleanum,* and *S. chacoense* under 14-hour light 10-hour dark (14/10) conditions. * Represents a two-tailed Welch’s t-test p-value of <0.05. As *S. candolleanum* did not tuberize during our experimental duration of 18 weeks, it was not included in this graph nor in the statistical analysis. B-J. The expression patterns of photoperiod-related tuberization genes genes in *S. tuberosum* (blue), *S. candolleanum* (purple), and *S. chacoense* (red) under long day conditions (mean, n = 2-3). *S. tuberosum* values are the mean ± standard deviation of all expressed alleles. Grey boxes represent nighttime conditions.

Changes in the phase of photoperiod-related genes are linked to differences in flowering time in Arabidopsis and other plant species (Yanovsky and Kay, 2002; Muranaka et al., 2022), therefore we analyzed their expression in our three species (Figs. 5B-J, S16). In potato, CO1 inhibits *SP5G* expression, a repressor of the tuberization promoter *SP6A* (Abelenda et al., 2016). The other two CO genes, *CO2* and *CO3,* are believed to play a similar role as *CO1*, but are more lowly expressed (Abelenda et al., 2016; Feke et al., 2025). We observed that under long days *S. chacoense* and *S. tuberosum CO2*, *CO3* and *SP5G*, but not *CO1,* display a later phase in comparison to *S. candolleanum* (Fig. 5B-F). *S. candolleanum SP6A* was not expressed under long days further supporting a very short critical photoperiod (Fig. 5G). Circadian input to this pathway is believed to be mediated via CDF1, a repressor of *CO1* and *CO2*, which is post-translationally regulated by FKF1 and GI (Kloosterman et al., 2013). We observed that in *S. candolleanum* the expression of *FKF1*, *GIa* and *GIb*, but not *CDF1,* was slightly advanced (Fig. 5E, H-J). These results suggest that changes in phase of photoperiod sensing genes might influence tuberization time in wild potatoes.

### Identification of potential causative genes using an *S. tuberosum*-*S. candolleanum* transcription factor network

Our results indicate that a significant portion of the genome and several physiological processes exhibit major delays in timing under diel conditions in domestic *S. tuberosum*. However, we have yet to determine the mechanism behind this delay. We hypothesized a cascading mechanism of transcriptional regulation between co-expression modules. To investigate this hypothesis, we generated a network representing potential regulatory relationships between the genes annotated with transcription factor function (TFs) found in our modules (Table S8, Table S9). Based on the known redundancy among TF protein family members, we collapsed TFs and their binding sites into what we termed “transcription factor groups” (TF groups) each containing TFs associated with one consensus motif (see Materials and Methods for details). Each "TF group" would in turn be regulated by motifs found in the promoters of the members of that group. We postulated that this simplified network would enable us to identify the principles of transcriptional regulation within the co-expressed modules.

We generated a network of 103 TF groups regulated by 45 motifs. Of the TF groups, 32 acted as both regulators and targets within our network (Fig. 6). While nearly two thirds of all motifs were only found in the promoters of five or fewer transcription factor groups, two motifs were each found to regulate over a quarter of the transcription factor groups within our network of motifs. Minimal overlap (less than 25%) was observed between the targets of these two motifs. Transcription factors regulating these two motifs represent potential hub genes for this network, and we focused on these moving forward.

**Figure 6.**
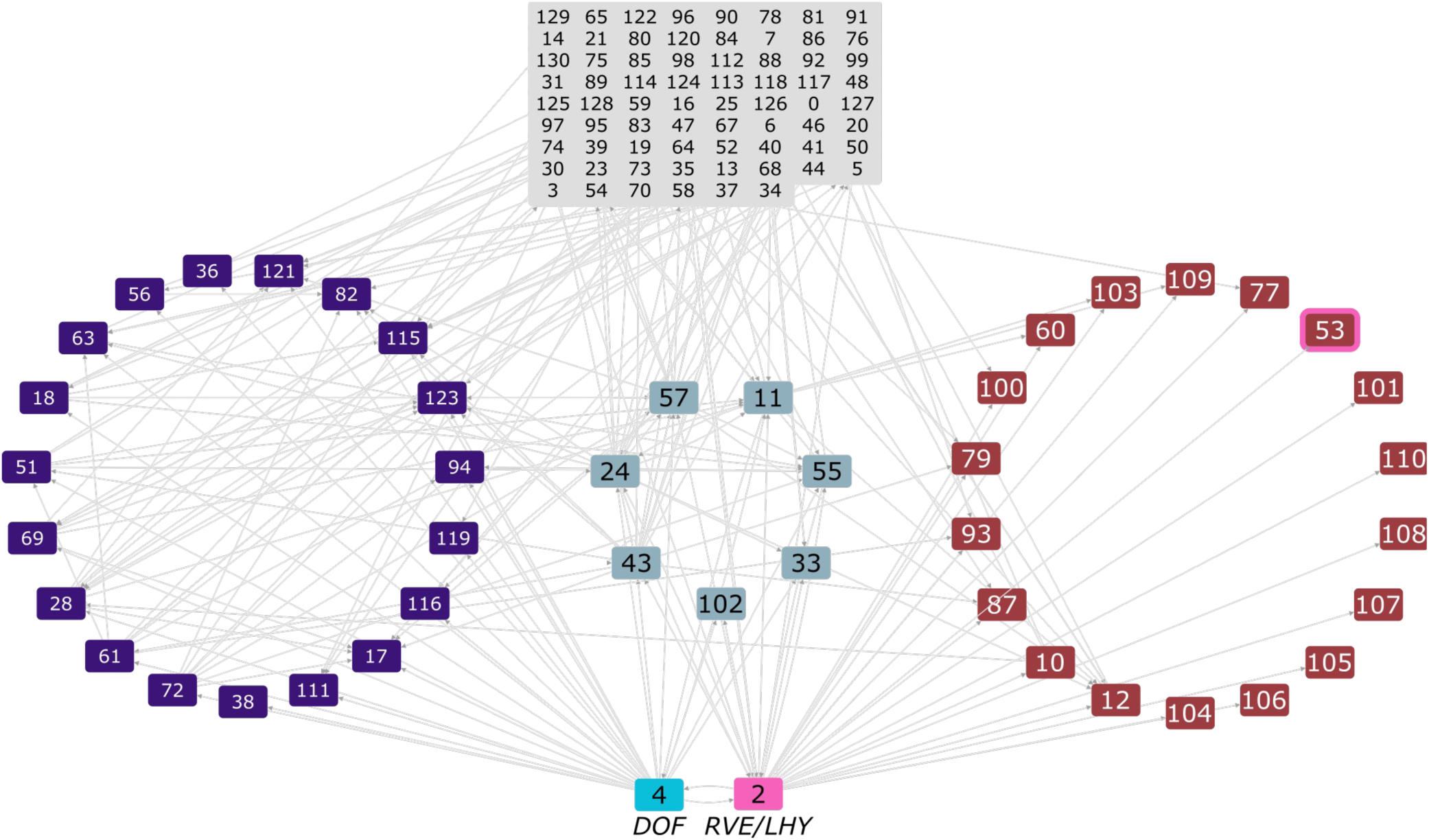
Diagram of the *S. candolleanum-S. tuberosum* transcription factor network. Individual boxes represent groups of TFs based on the similarity of their binding sites in Arabidopsis and are labeled with the TF group number. The cyan box (group 4) contains the *CDF*s, while the pink box (group 2) contains the *RVE*s and *LHY*. Purple boxes are those that are targeted by the CDFs, red boxes are those that are targeted by the RVEs, and the blue-grey boxes are those that are targeted by both the RVEs and the CDFs. Grey boxes represent other transcription factors within our network that are not directly targeted by either the RVEs or the CDFs. The red box with a pink outline contains *DEL1*.

The motif that controlled the greatest number of TF groups in our network was WAAAG, associated with ’Group 4’ transcription factors. This motif is associated with the Cycling DOF Factor genes (CDF), which have been implicated in photoperiodic reproductive processes such as flowering and tuberization (Imaizumi et al., 2005; Kloosterman et al., 2013). None of the syntelogs within Group 4 exhibited significant phase delays in *S. tuberosum* in comparison to *S. candolleanum* (Fig. S17). However, *S. tuberosum* cv. Atlantic encodes two truncated alleles of *CDF1* that are unable to interact with the CDF-degrading protein FKF1 and thus are believed to be constitutively expressed at the protein level (Kloosterman et al., 2013). To test whether *CDF1* could be affecting the timing of expression in *S. tuberosum*, we investigated whether genes differentially expressed in *CDF1*-misexpression lines (Shaikh et al., 2025) were present in our modules. We found that approximately 47% of the 2,560 syntelogs differentially expressed in either truncated *CDF1*-overexpression or *CDF1*-RNAi lines were found in our modules. In contrast, only 9% of syntelogs are misregulated in these lines genome-wide, indicating that a significant portion of CDF1-regulated genes displayed diel regulation (Fig. 7A). We found enrichment for genes that are impacted by *CDF1* misexpression in 10 of our 18 modules when compared to all expressed syntelogs, including module 16, wherein nearly 40% of genes were differentially expressed (Fig. 7B). This suggests that *CDF* genes may play a significant role in the gene expression patterns we observed within some of our modules.

**Figure 7.**
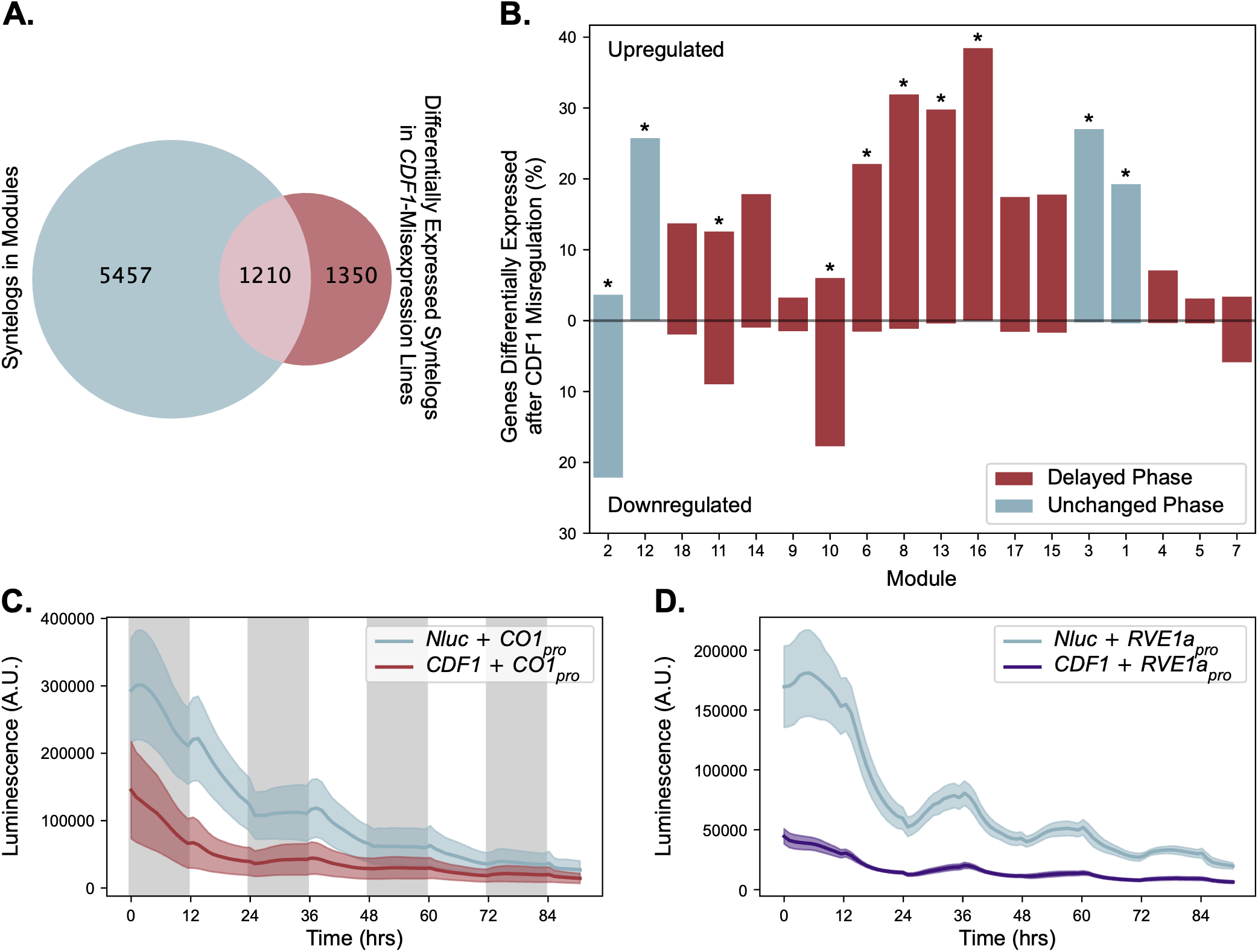
*CDF1* has a significant impact on rhythmic gene expression. A. Venn diagram of syntelogs within modules compared to those that were differentially expressed when *CDF1* is mis-expressed. B. The proportion of genes in each module that is differentially expressed when *CDF1* is misregulated. The total bar height represents the total number of genes that were differentially expressed, while the bar height above the central line represents those that were upregulated and the bar height below the central line represents those that were downregulated. Modules are ordered by the time of maximum expression, from the earliest to the latest. Blue modules are those that exhibited no phase difference between *S. tuberosum* and *S. candolleanum,* while red modules were those that are delayed in *S. tuberosum.* Statistical enrichment was determined using a Fisher’s Exact test with all expressed syntelogs used as the background, * represents a Benjamini-Hochberg-corrected *p*-value <0.05. C.-D. *CDF1* acts as a direct repressor of C. *CO1* and D. *RVE1a*. Either *CDF1* (red or purple) *or Nluc* (blue) was co-infiltrated with firefly luciferase driven by either A) the *CO1* promoter (red) or B) the *RVE1a* promoter (purple). Luminescence was measured approximately hourly, starting at ZT12.

The other motif, AAATATCT, was associated with Group 2 TFs. This motif, known as the evening element, is a well characterized motif involved in circadian clock regulation and the known binding site of several core clock transcription factors including At*LHY* and the *AtRVEs* (Alabadi et al., 2001; Rawat et al., 2009; Rawat et al., 2011). While four of the five potato genes in our dataset that were hypothesized to bind to the evening element exhibited phase advances in *S. tuberosum*, the fifth, *RVE1a*, had a significant delay in *S. tuberosum* compared to *S. candolleanum* (Figs. 2A, S4). Interestingly, we found that *RVE1a* contained both the evening element and the CDF motif in its promoter in *S. candolleanum* and all four *S. tuberosum* alleles. In contrast, none of the other *RVE* or *LHY* genes contained the evening element in their promoters, nor was the CDF motif consistently found in all alleles of these genes in both species (Fig. S18). This suggests that the evening element, and *RVE1a* in particular, may serve as a potential hub of our network, receiving inputs from the clock and from CDF1, thus leading to the phase delay in *S. tuberosum* due to the constitutive protein expression of CDF1.

If the *RVE*/*LHY* genes are key factors for the global phase delay observed in *S. tuberosum*, then we would hypothesize that the evening element would be enriched in the promoters of at least one of the delayed modules. We found that the evening element was the most strongly enriched element in three modules, modules 1, 3, and 6 (Fig. 8). Only one of these modules, module 6, exhibited a phase delay in *S. tuberosum*. This is not surprising, since we expect that the combinatorial effect of multiple transcription factors determines the phase of expression and multiple factors can bind to the evening element (Rawat et al., 2011). We observed that the location of the evening element with respect to the transcriptional start site differs between the unshifted modules 1 and 3 and the shifted module 6 (Fig. S19), indicating that they might be regulated by different sets of TFs.

**Figure 8.**
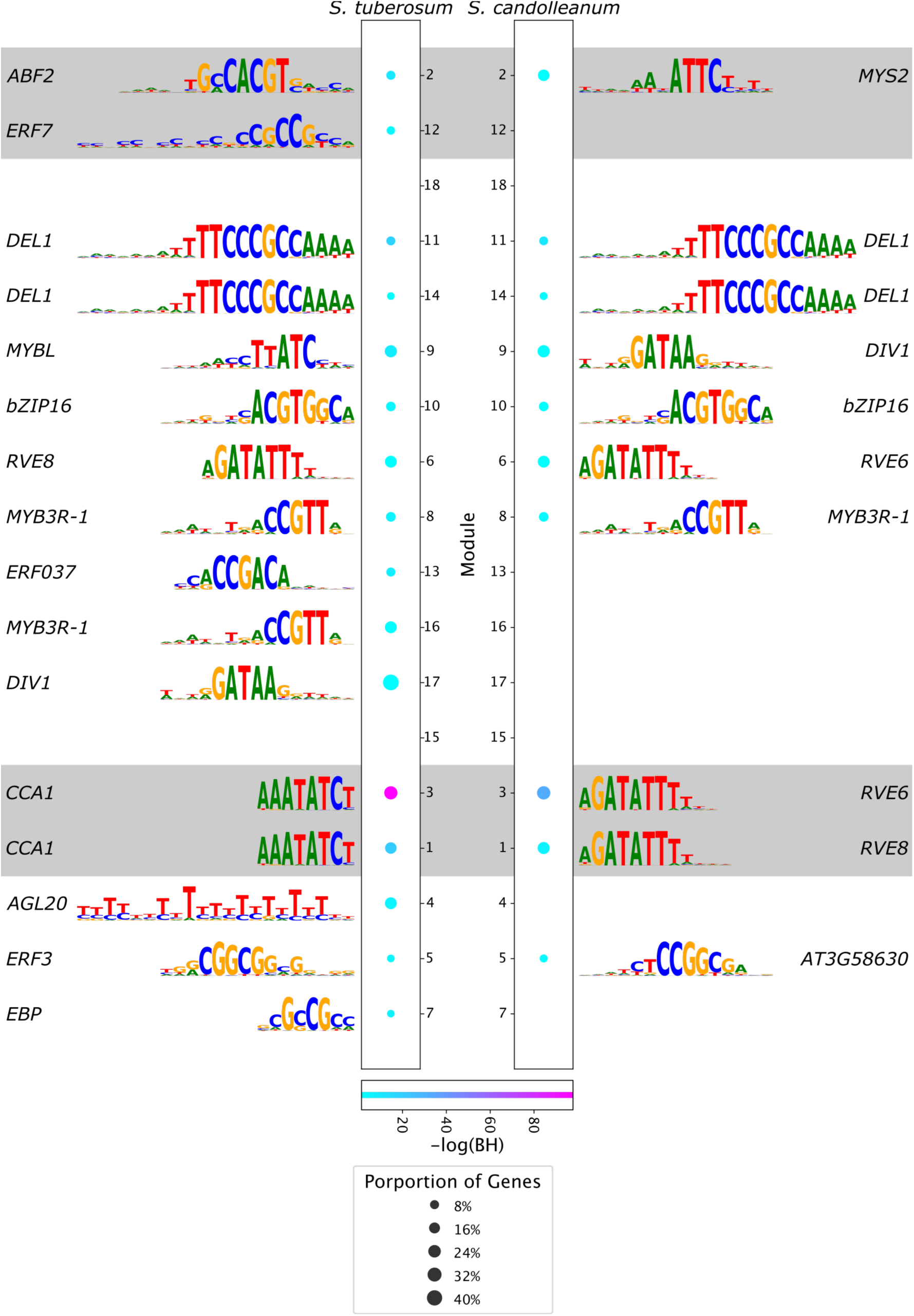
The most highly enriched motif in *S. tuberosum* (left) and *S. candolleanum* (right) within the promoters of each module. Sequence logos of each motif are provided to the left and right, while the central plot represents the proportion of genes within each module that possessed that motif within its promotors (circle size) and the p-value of the enrichment of that motif (circle color). Modules are ordered from earliest timepoint of maximum expression to latest. Modules with shaded backgrounds are those that exhibited no phase difference between *S. tuberosum* and *S. candolleanum,* while unshaded modules were those that are delayed in *S. tuberosum.* Gene names displayed to the left and right of each sequence logo are the Arabidopsis gene associated with each motif.

Since we do not find the evening element enriched in the promoters of additional shifted modules, we next queried whether transcription factors in our shifted module 6 could be regulating other shifted modules. To identify transcription factors that could link transcriptional regulation between modules we searched for transcription factors in module 6 that contained an evening element in their promoters. One of these transcription factors is DP-E2F-like (DEL1). DEL1 binds to the TTCCCGCCAAAA motif, which was found to be the most significantly enriched motif in the promoters of genes in modules 11 and 14 in both *S. tuberosum* and *S. candolleanum* (Fig. 8). *DEL1* has been implicated in the regulation of DNA replication and the endocycle (Vlieghe et al., 2005; Lammens et al., 2008; Heyman et al., 2017). Correspondingly, we found functional enrichment of genes relating to DNA replication and wrapping within modules 11 and 14. Notably, approximately 45% of the genes in module 11 with the *DEL1* motif in their promoters were associated with DNA replication, while more than 84% of genes associated with DNA replication in module 11 possessed the *DEL1* motif in their promoters (Table S10).

### CDF1 represses the activity of the *RVE1a* promoter

Our transcriptional network analyses indicate that CDF1 might be regulating *RVE1a* expression in potato. Previous DAP-seq studies have shown that CDF1 is able to bind to the *RVE1a* promoter *in vitro* (Shaikh et al., 2025), suggesting the presence of a direct interaction. We tested whether the truncated CDF1 from *S. tuberosum* was able to influence *RVE1a* promoter transcription. As previously reported, *S. tuberosum* CDF1 represses the transcriptional activity of the *CO1* promoter (Fig. 7C) (Kloosterman et al., 2013). The presence of CDF1 also repressed the activity of the *RVE1a* promoter in a transient transformation assay (Fig. 7D), suggesting that CDF1 is a transcriptional repressor of *RVE1a* in potato.

## Discussion

The variation of circadian period among wild potatoes supported the hypothesis that a long period helps plant survival under the long summer days found at higher latitudes by causing delays in phase (Müller et al., 2016; Hardigan et al., 2017). However, the lack of variation among potato cultivars raised the question of whether those delays are important in agricultural settings. Here, we present a study on the effects of natural differences in the circadian period on gene expression patterns genome-wide using three tuber-bearing *Solanum* species, two wild species and a domesticated *S. tuberosum* cultivar. By including both wild and domestic varieties, we were able to discover a clock independent mechanism leading to delayed phase in modern cultivated potato.

It has long been hypothesized that varying the circadian period will lead to alterations in the phase of clock-regulated genes under diel conditions (Aschoff and Pohl, 1978). Dawn, acting as the zeitgeber, will reset the clock daily, and thus alterations in period will act to delay or advance the timing of gene expression relative to dawn. In all three species there was a tight correlation between the period and the diel phase of orthologs to well-characterized circadian clock genes. The short-period *S. candolleanum* and *S. tuberosum* exhibited significant earlier phases than the long-period *S. chacoense* (Fig. 2A). This suggests that regarding the core clock genes themselves, our understanding of the connection between circadian period and phase is well-supported by gene-expression data from both wild and domesticated species. However, only within the wild species did the average genome-wide diel phase correlate with the circadian period. In contrast, we observed significant delays in gene expression in the modern cultivated potato *S. tuberosum* despite their fast free running period. Cultivated tomato are characterized for having both late phase and a long period of free running leaf movement rhythms that appeared as a two-step process during domestication (Müller et al., 2016; Muller et al., 2018). These modifications also lead to delayed phase of gene expression under diel conditions in modern cultivated tomatoes (Müller et al., 2016) supporting the idea that a delayed phase plays a key role in plant productivity under higher latitudes.

The physiological implications of delayed phase under light/dark conditions remain unclear. We measured physiological rhythms of processes that have been shown to be under circadian control in other species, such as light reactions of photosynthesis (Litthauer et al., 2015), starch and sugar content (Lu et al., 2005; Graf et al., 2010) and leaf movement (Nozue et al., 2011). Only leaf movement displayed significant delayed phases in *S. chacoense* and *S. tuberosum* correlating with delays in gene expression (Fig. 4). The function of these hyponasty rhythms could be to avoid shading by neighboring plants and/or protect leaves from stressors such as high light and temperature occurring at the second time of the day (Woodley Of Menie et al., 2019; Havko et al., 2020). The combination of phase changes of photosynthesis gene expression and leaf movement rhythms could help optimize light capture during the day, leading to similar phases of photosynthetic traits in all three genotypes (Figs. S10, S11).

In Arabidopsis, the circadian clock has been proposed to help optimize growth by regulating transitory leaf starch levels (Graf et al., 2010). We observed that diel oscillations of sucrose content were delayed in cultivated potato, indicating that the timing of metabolism is affected by the delays in gene expression. However, diel starch rhythms varied strongly between the three species. Notably, the short period *S. candolleanum* was the only species with decreased starch content under long days, suggesting that early phases could be detrimental under very long photoperiods. In Arabidopsis, extreme long photoperiods also lead to a decrease in transitory leaf starch (Sulpice et al., 2014). In contrast, *S. chacoense* displayed extremely high starch levels with no detectable oscillations under long days. This is similar to what has been observed in some understory Ginseng species under elevated light intensities (Miskell et al., 2002), indicating that plants have different strategies to optimize leaf carbohydrate metabolism.

Variation of the circadian period and/or diel phase has been associated with differences in photoperiod sensitivity in wild and domesticated species (Salmela et al., 2016; Greenham et al., 2017; Rees et al., 2021; Muranaka et al., 2022; Wittern et al., 2023; Helmsorig et al., 2024). A strong decrease in photoperiod sensitivity in cultivated potato is caused by truncated *CDF1* alleles (Kloosterman et al., 2013). However, we also observe significant differences in tuberization time between our short and long period accessions which both encode full length CDF1 versions (Fig. 5A). The current model of tuberization time in potato contains similar mechanisms of regulation as flowering time in Arabidopsis, including the circadian and light regulation of a light-stabilized protein (Abelenda et al., 2014). Further studies are needed to determine whether phase changes modulate daylength sensitivity in potato.

Our transcription factor network enabled the identification of two cis-regulatory elements which may act as hubs for the domestication-induced delay in phase in cultivated potato. We hypothesize that these elements mediate regulation by CDF1 and RVE1a. The truncated versions of *CDF1* found in cultivated potato lead to constitutive levels of this protein yet remain transcriptionally active (Kloosterman et al., 2013). In Arabidopsis, the lack of AtCDF1 post-translational regulation by FKF1 also leads to a delayed phase (Nelson et al., 2000), suggesting conserved roles of CDF proteins. In potato, truncated *CDF1* forms, including the *CDF1.2* allele found in *S. tuberosum* cv. Atlantic, have been associated with multiple physiological effects in addition to tuberization time, including nitrogen metabolism, plant senescence, and increased vigor (Shi et al., 2024; Amundson et al., 2025; Shaikh et al., 2025). We hypothesize that CDF1 misexpression triggers a cascading mechanism leading to delayed expression of a large percentage of the rhythmic genome, at least partially by regulating *RVE1a*. In Arabidopsis, *RVE1* influences hypocotyl growth but not circadian rhythms (Rawat et al., 2009). Our analyses indicate that in potato *RVE1a* could be regulating growth via *DEL1*, which, in Arabidopsis has been associated with cell growth (Chandran et al., 2014; Nakagami et al., 2020).

Our findings show that changes in phase are associated with crop productivity at higher latitudes. Although phase differences can be generated through altering the circadian period, as was observed in our two wild species, alternative mechanisms that change phase but not the circadian period also have significant physiological effects, as was observed in the domestic cultivar. Understanding the mechanisms of how alterations in *CDF1* lead to differing plant vigor and tuberization phenotypes will shed light on how this critical potato domestication gene is able to coordinate these responses.

## Methods

### Plant materials and growth conditions

*S. tuberosum* cv. Atlantic, *S. candolleanum* (PI 210044) and *S. chacoense* "M6" (Jansky et al., 2014) were grown from stem cuttings as described previously (Feke et al., 2025). For transcriptome analyses, three and five terminal leaflets were collected from the third and fourth newly expanded leaf for *S. chacoense* and *S. candolleanum*, respectively. *S. tuberosum* tissue was collected as described previously (Feke et al., 2025). For short-day (12 hour light/12 hour dark, 22°C day, 18°C night) experiments, *S. tuberosum* cv. Atlantic plants were nine weeks old, *S. chacoense* plants were seven weeks old, and *S. candolleanum* plants were eleven weeks old at the time of harvest, and all plants had tubers. For long-day (16 hour light/8 hour dark, 22°C day, 18°C night) experiments, all genotypes were 4.5 weeks old at the time of harvest. For biomass measurements all samples were harvested 5 weeks after planting. For tuberization measurements, plants were grown at the indicated photoperiods (see figure legends) and checked for tuber formation weekly; the experiment was stopped after 14 weeks.

### Genome sequencing, assembly, and annotation

See Supplemental File 1 for an extended methods description of the methodology of the genome sequencing, assembly, and annotation.

### Transcriptome analysis and generation of the gene regulatory network

See Supplemental File 1 for an extended methods description of the transcriptomic analysis. In brief, JTK was used to determine rhythmicity according to a *p*-value<0.001 (Hughes et al., 2010). Syntelogs were considered rhythmic in *S. tuberosum* when at least one allele was rhythmic. Functional annotation was based on the Mapman4 terms (Bolger et al., 2021), using the high confidence representative gene models, and was the same for all genes belonging to a syntelog (Datasets S14-16). *S. tuberosum* cv. Atlantic mapman annotation has been previously reported (Feke et al., 2025). Mapman terms were considered significantly enriched if they displayed a Holm-corrected *p*-value < 0.05 and the fraction of genes in the study sample assigned to that term was at least 25% greater than the fraction of genes in the population with that term. Co-expression modules were generated based on concatenated *S. tuberosum* and *S. candolleanum z*-scored expression patterns for each S*. tuberosum* allele based on syntelog identity. The gene-regulatory network was generated based on the annotated transcription factor binding site for the *A. thaliana* gene orthologous to each syntelog, with all syntelogs with similar binding motifs collapsed into a single TF group (Jin et al., 2017)(Table S8). The presence of a transcription factor binding site within promoters was identified using Hypergeometric Optimization of Motif EnRichment (HOMER) (Heinz et al., 2010).

### Measurement of circadian period

Circadian period was measured using delayed fluorescence as described previously (Hardigan et al., 2017).

### Physiological Assays

See Supplemental File 1 for an extended methods description of the metabolite and photosynthesis assays.

### Transient Promoter Assays

*pMDC84-CDF1*, promoter reporter constructs *pCO1pr:LUC* and *pRVE1apr:LUC,* empty vector control *pMDC84-NLUC*, and *pCB-19HA* silencing suppressor were transformed into *Agrobacterium tumefaciens* (GV3101). Please see Supplemental File 1 for cloning details. Overnight *A. tumefaciens* cultures were resuspended 10 mM MES, pH 5.8, 10 mM MgSO_4_ and 225 μM acetosyringone to 0.5 OD_600_. Four-week-old *N. benthamiana* plants were grown under 12 hour light/12 hour dark, 350 mmol m^-2^ s^-1^ white light. *A. tumefaciens* suspensions were mixed using a 1:2:1 reporter: transcriptional regulator: p19 silencing suppressor ratios. Each half of each leaf was infiltrated using the transcription regulator and the reporter gene or the NLUC control and the reporter gene. Three days after transformation, 0.6 cm punches were cut out of the infiltrated regions and transferred to a 96 well plate containing 100 mL 1/2 concentration liquid Murashigue & Skoog media supplemented with 40 mg ml^−1^ D-luciferin (Gold Biotechnology). Luminescence was measured as described previously (Liu et al., 2016) under light/dark cycles of 25 mmol m^-2^ s^-1^ fluorescent white light.

### Figure generation and data analysis dependencies

Figures were generated using the python packages matplotlib (Hunter, 2007), seaborn (Waskom, 2021), matplotlib-venn (https://github.com/konstantint/matplotlib-venn), and seqlogo (github.com/betteridiot/seqlogo). Statistical tests were implemented as described in the respective figure legends. Additional packages not cited elsewhere that were integral to the data analysis include pandas (The pandas development team), numpy (Harris et al., 2020), and SciPy (Virtanen et al., 2020).

## Data availability

All sequencing data can be accessed under the NCBI Sequence Read Archive BioProjects PRJNA826514, PRJNA826516, PRJNA957457, and PRJNA1093480. The code used in this study is available at GitHub (TBD). The large datasets (Datasets S1-S15) have been deposited in the Dryad repository (TBD). Genomic resources for *S. chacoense* and *S. candolleanum* can be accessed via SpudBD (https://spuddb.uga.edu/index.shtml).

## Acknowledgements

We thank Shannon Donnelly, Berkley Walker and Donghee Hoh for useful advice and discussions on the photosynthesis measurements. We are grateful to Linsey Newton for the generation of the *pGreenII-0800-luc-GW* construct, and to Marisa Laitinen for her assistance in collecting time course tissue samples.

## Author contributions

C.R.B and E.M.F conceived the research. A.F., B.V., K.A., J.C.W, S.H., C.R.B., E.M.F., designed and carried out the experiments. A.F., B.V, J.P.H., S.H., J.C.W., D.H., and E.M.F. performed the data analyses. A.F. and E.M.F. wrote the manuscript. All authors revised and edited the manuscript.

## Supplementary material

Supplemental file 1. Extended methods description.

## Supplemental figures

**Figure S1.**
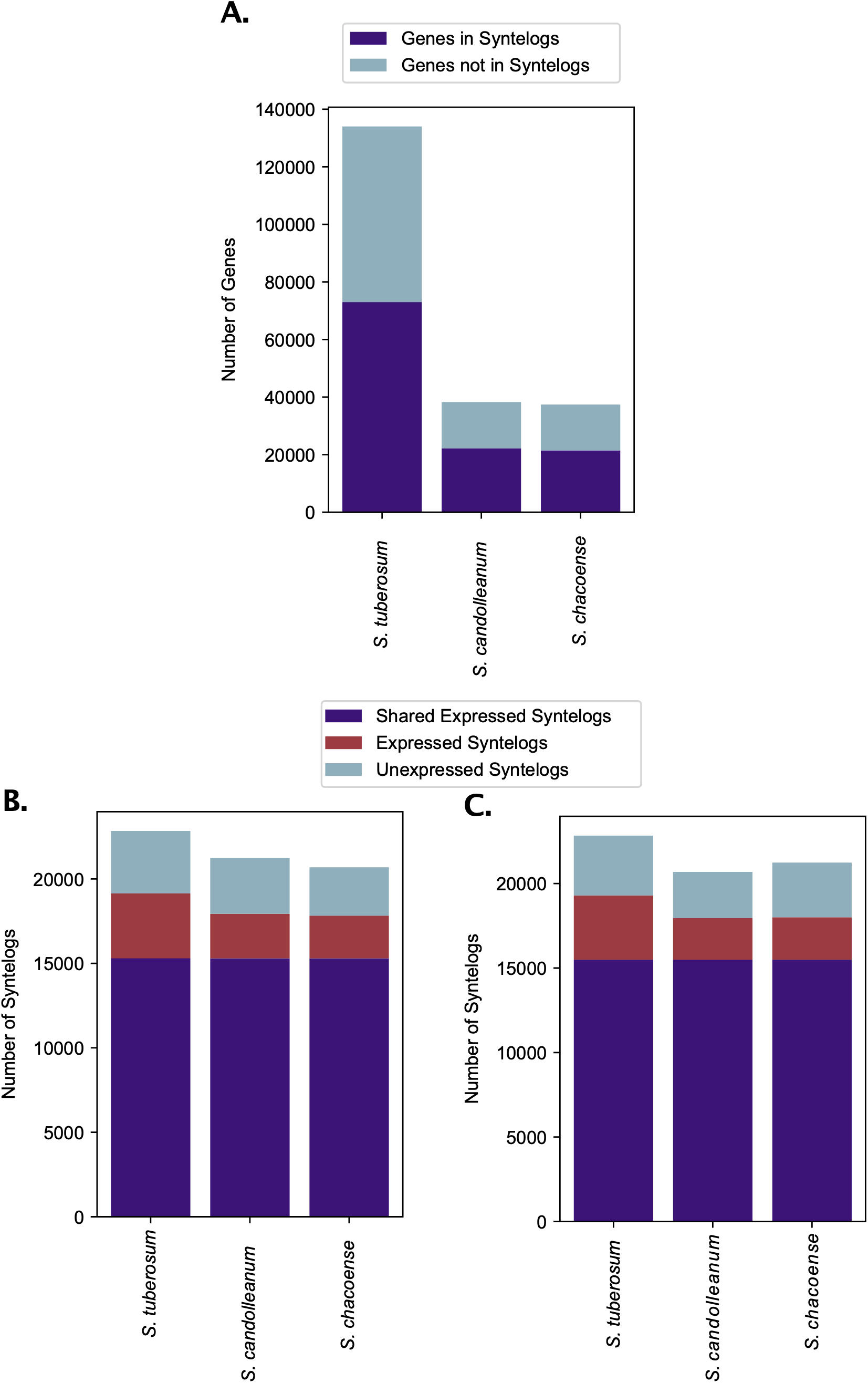
Syntelog coverage of the three *Solanum* species. A. The proportion of genes within potato genomes that are captured by our stringent syntelog definition. B-C. The proportion of syntelogs that are expressed in *S. tuberosum*, *S. candolleanum, and S. chacoense* under B. short day and C. long day conditions. Unexpressed syntelogs are those that are not expressed in any species, shared expressed syntelogs are those that are expressed in all three species, and expressed syntelogs are those that are expressed in that species but not all three.

**Figure S2.**
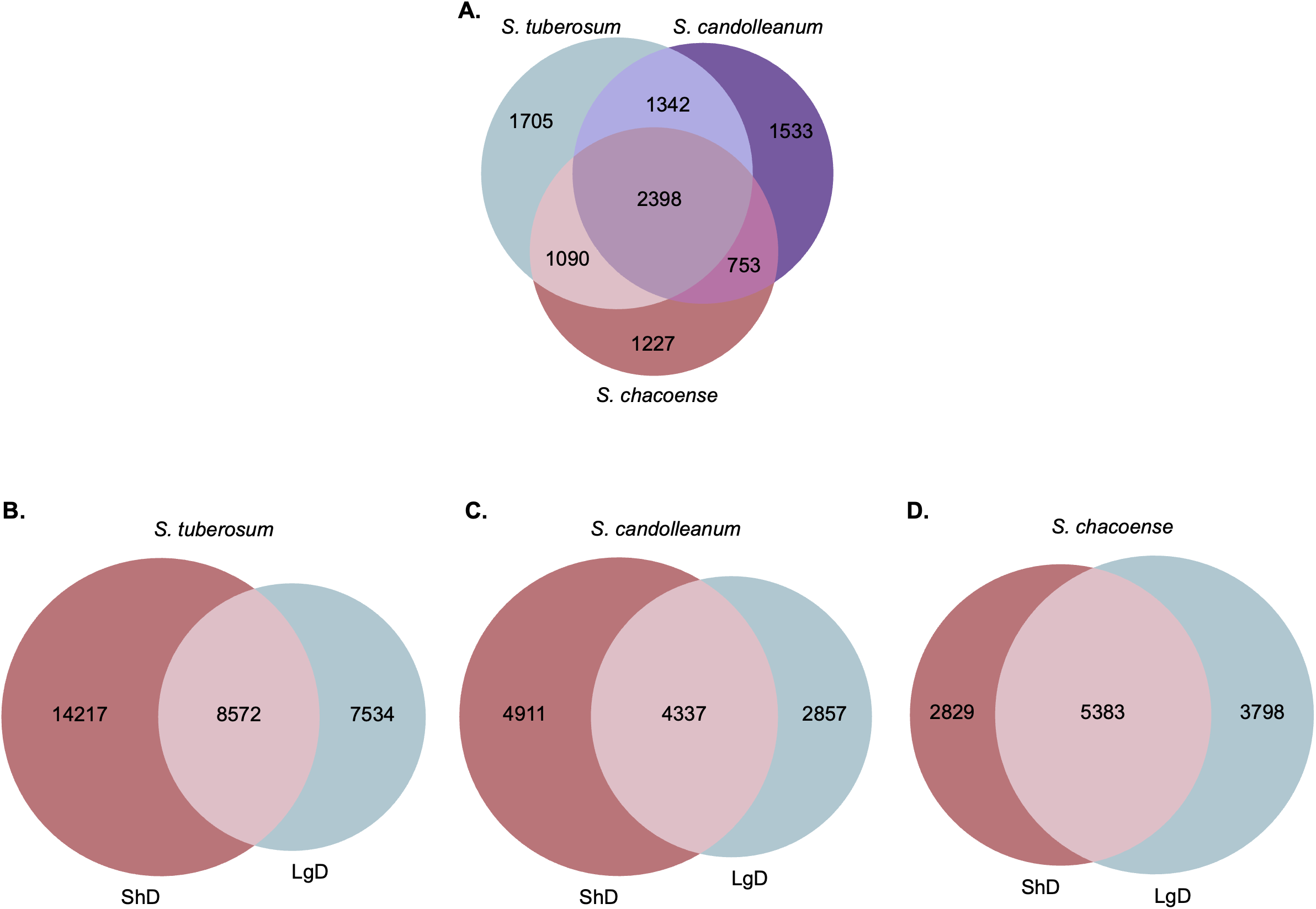
Overlap of rhythmic gene expression between experiments. A. Venn diagram of rhythmic genes in all 3 species under short day conditions. B-D. Venn diagram of rhythmic genes in both photoperiods in B. *S. tuberosum* C. *S. candolleanum,* and D. *S. chacoense*.

**Figure S3.**
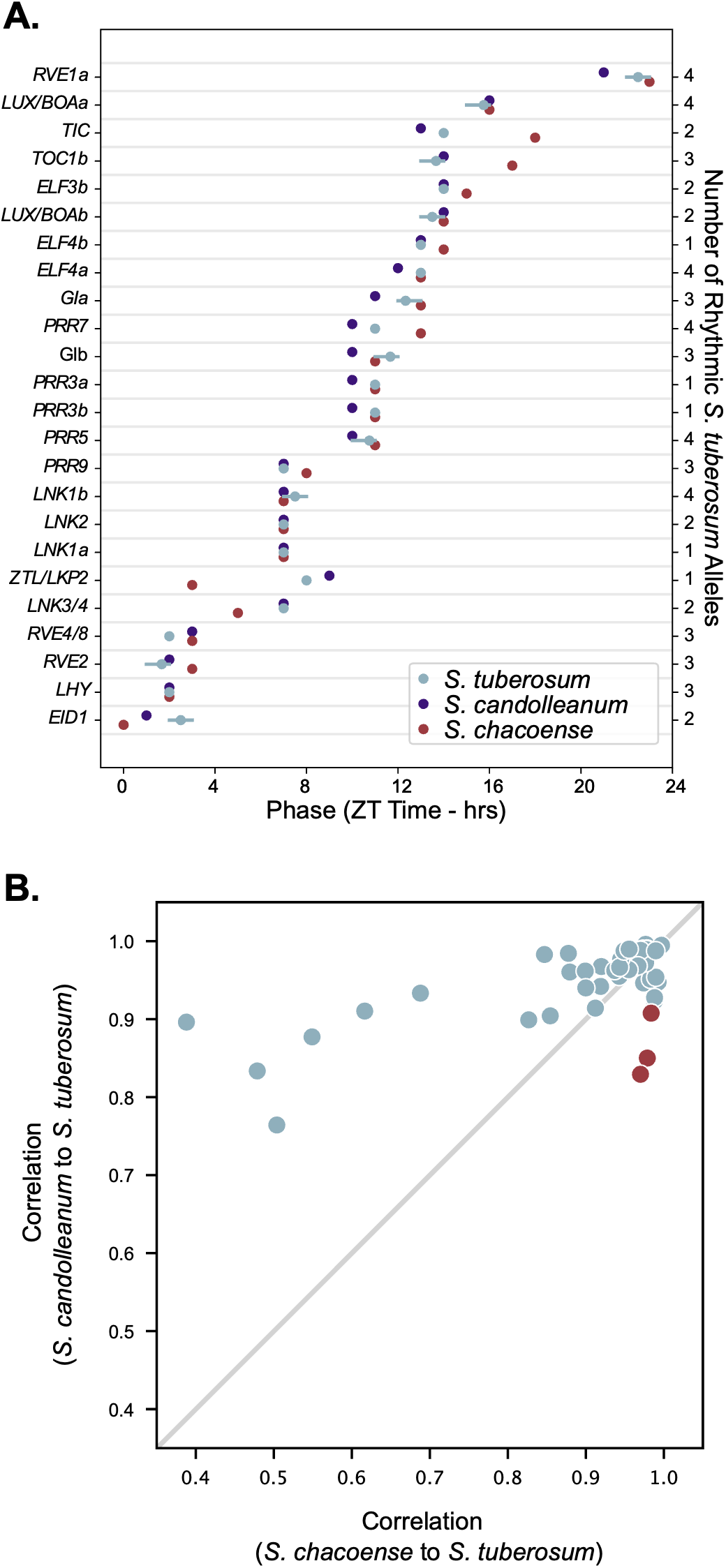
Circadian period correlates with phase under long day conditions. A. The phase of expression of rhythmic clock genes in *S. tuberosum* (blue), *S. candolleanum* (purple), and *S. chacoense* (red) under long day conditions. As the *S. tuberosum* assembly was haplotype-resolved, anywhere from 1-4 alleles can be included for each *S. tuberosum* datapoint. For this species, the circles represent the mean phase of all expressed and rhythmic alleles, while the horizontal lines represent the range of phases for each gene. The number of alleles that is included for each gene is listed on the right y-axis. B. Pearson correlations between the gene expression patterns of rhythmic clock genes *S. tuberosum* compared to either wild species under long day conditions. *RVE1a* is highlighted in red.

**Figure S4.**
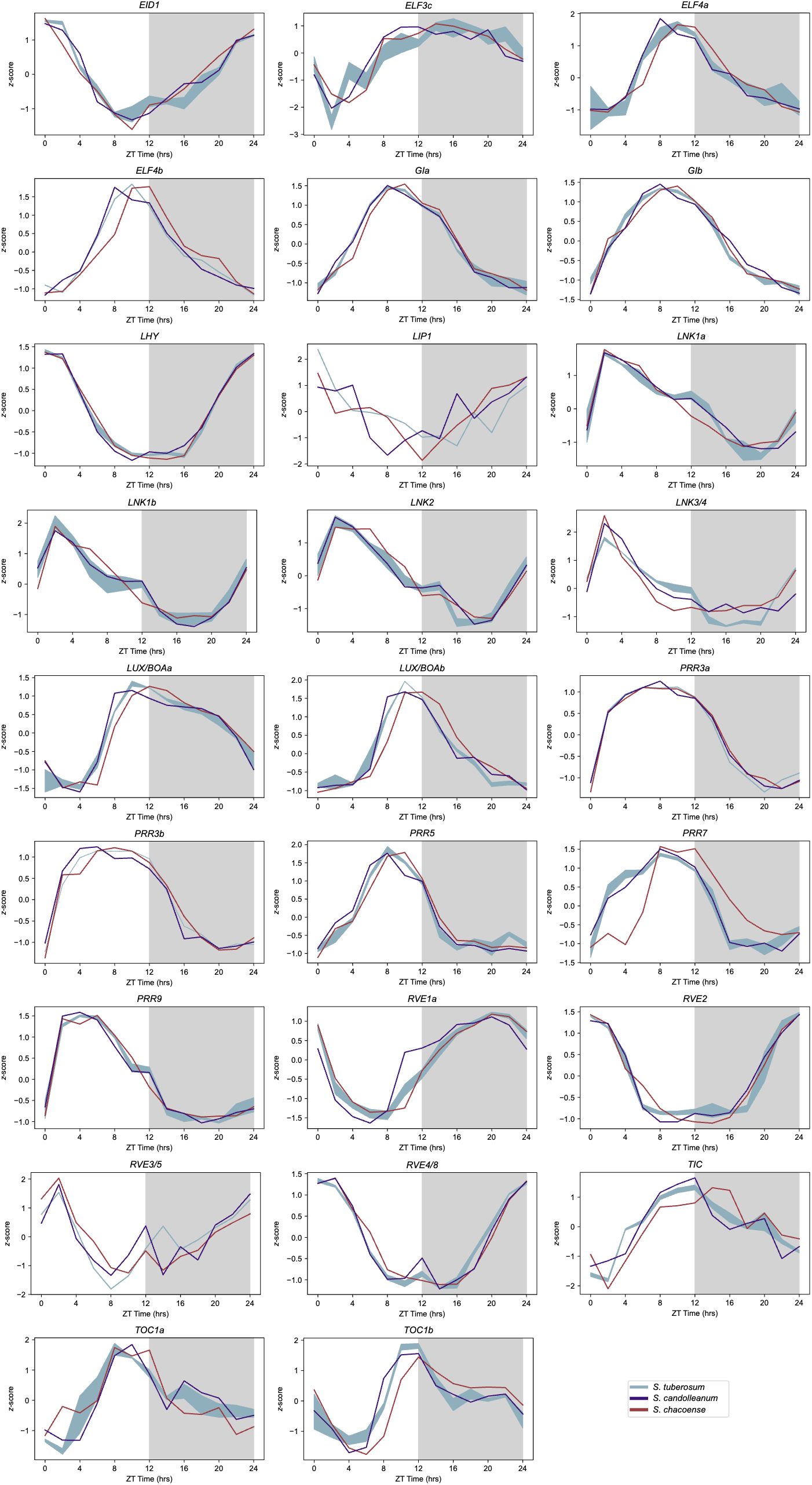
The expression patterns of rhythmic clock genes in *S. tuberosum* (blue), *S. candolleanum* (purple), and *S. chacoense* (red) under short day conditions. *S. tuberosum* values are the mean ± standard deviation of all rhythmic alleles. Grey boxes represent nighttime conditions.

**Figure S5.**
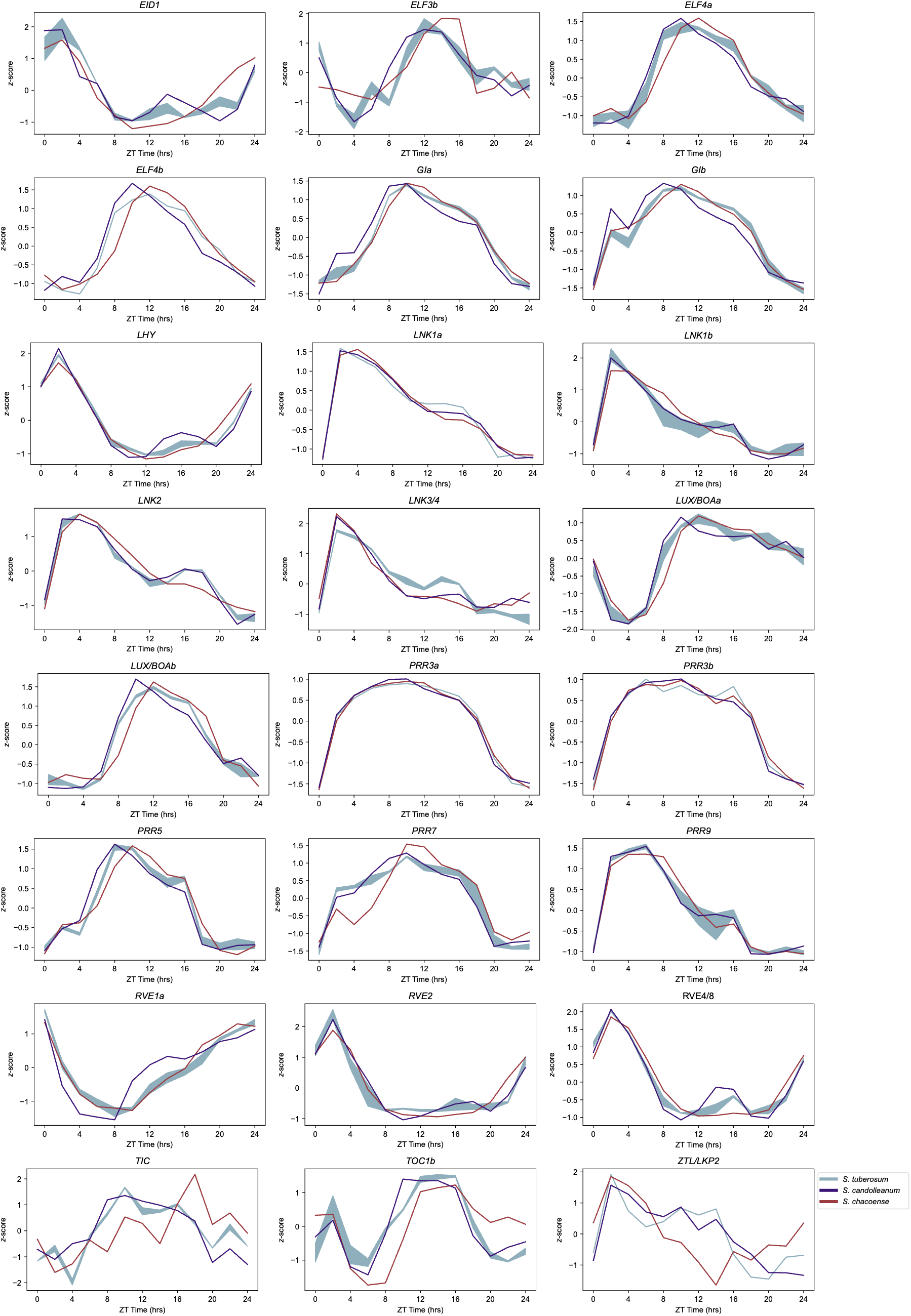
The expression patterns of rhythmic clock genes in *S. tuberosum* (blue), *S. candolleanum* (purple), and *S. chacoense* (red) under long day conditions. *S. tuberosum* values are the mean ± standard deviation of all rhythmic alleles. Grey boxes represent nighttime conditions.

**Figure S6.**
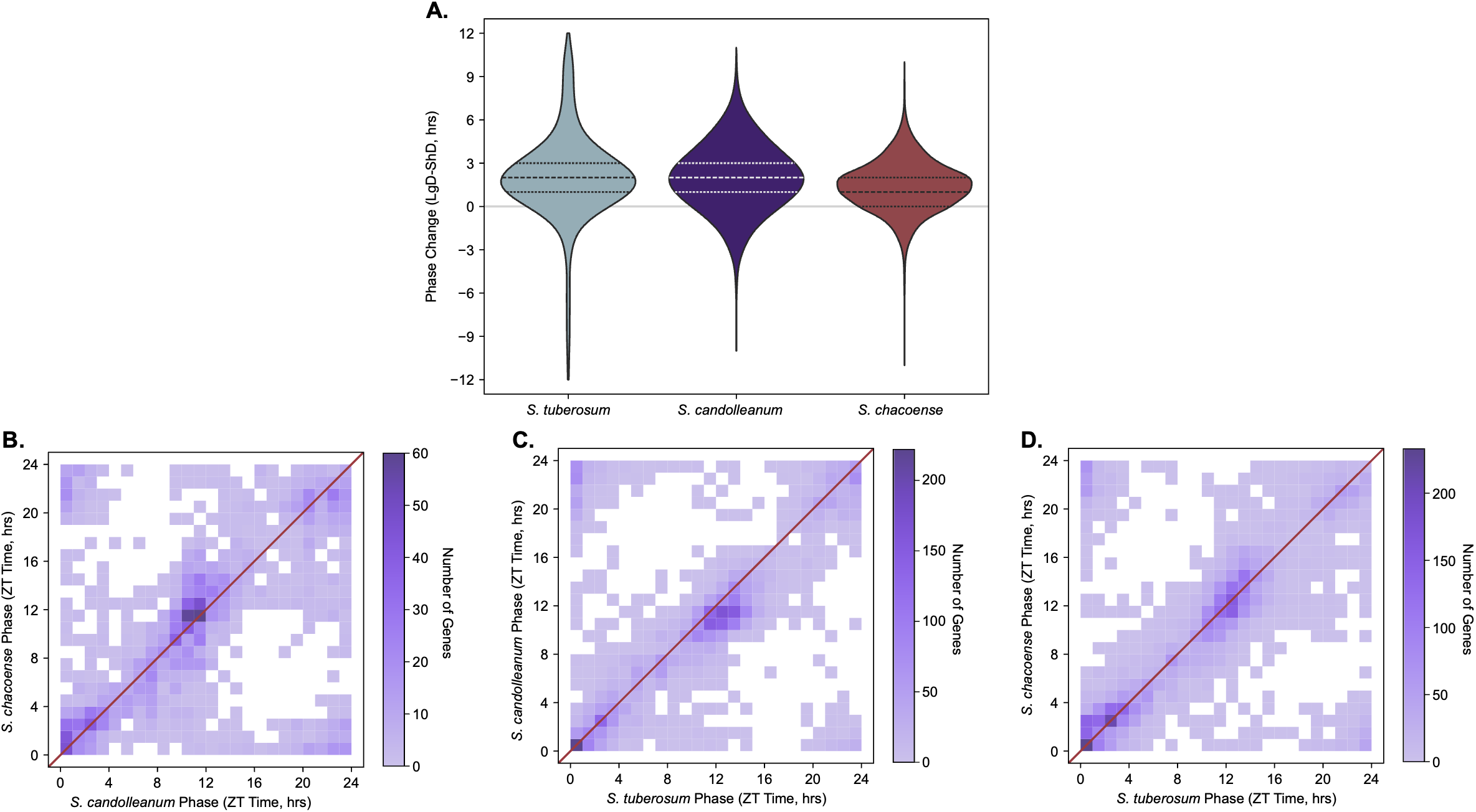
Phase delays are maintained under long day conditions. A. Violin plot of phase changes with photoperiod in *S. tuberosum*, *S. candolleanum,* and *S. chacoense.* Positive values represent those where phase is later in long days than in short days, while negative values represent those where phase is later in short days than in long days. Dashed lines represent the median, while the dotted lines represent the first and third quartiles. B-D Two-way histograms of phase across all genes that were rhythmic under short day conditions in both B. *S. candolleanum* and *S. chacoense* C. *S. candolleanum* and *S. tuberosum*, or D. *S. chacoense* and *S. tuberosum.* The red lines represent the points where phase is equal in both species. *S. tuberosum* is significantly delayed in comparison to *S. candolleanum* (Wilcoxon Signed Rank Test *p*-value<0.001), while both species exhibit no significant differences when compared to *S. candolleanum* (Wilcoxon Signed Rank Test *p*-value>0.05)

**Figure S7.**
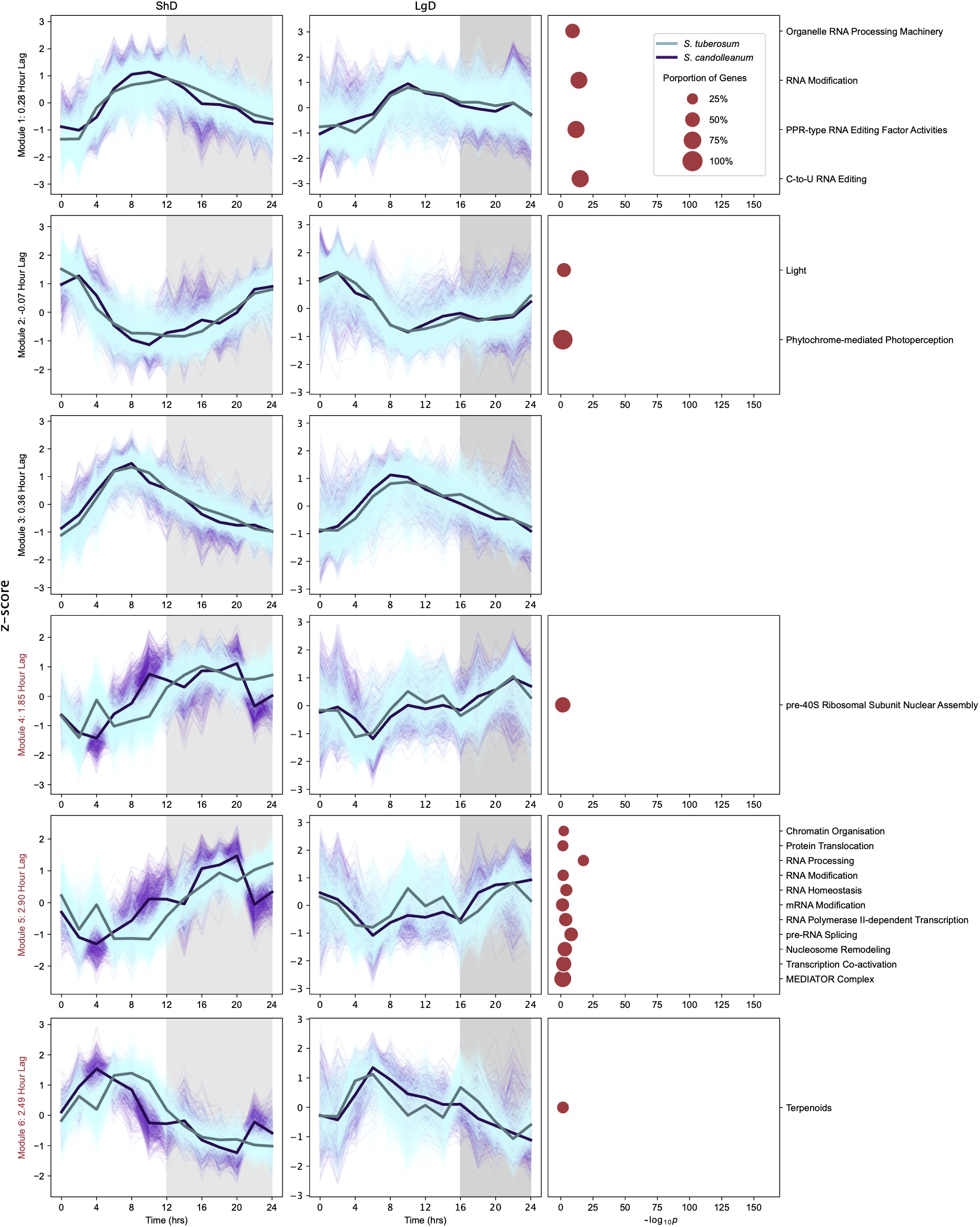
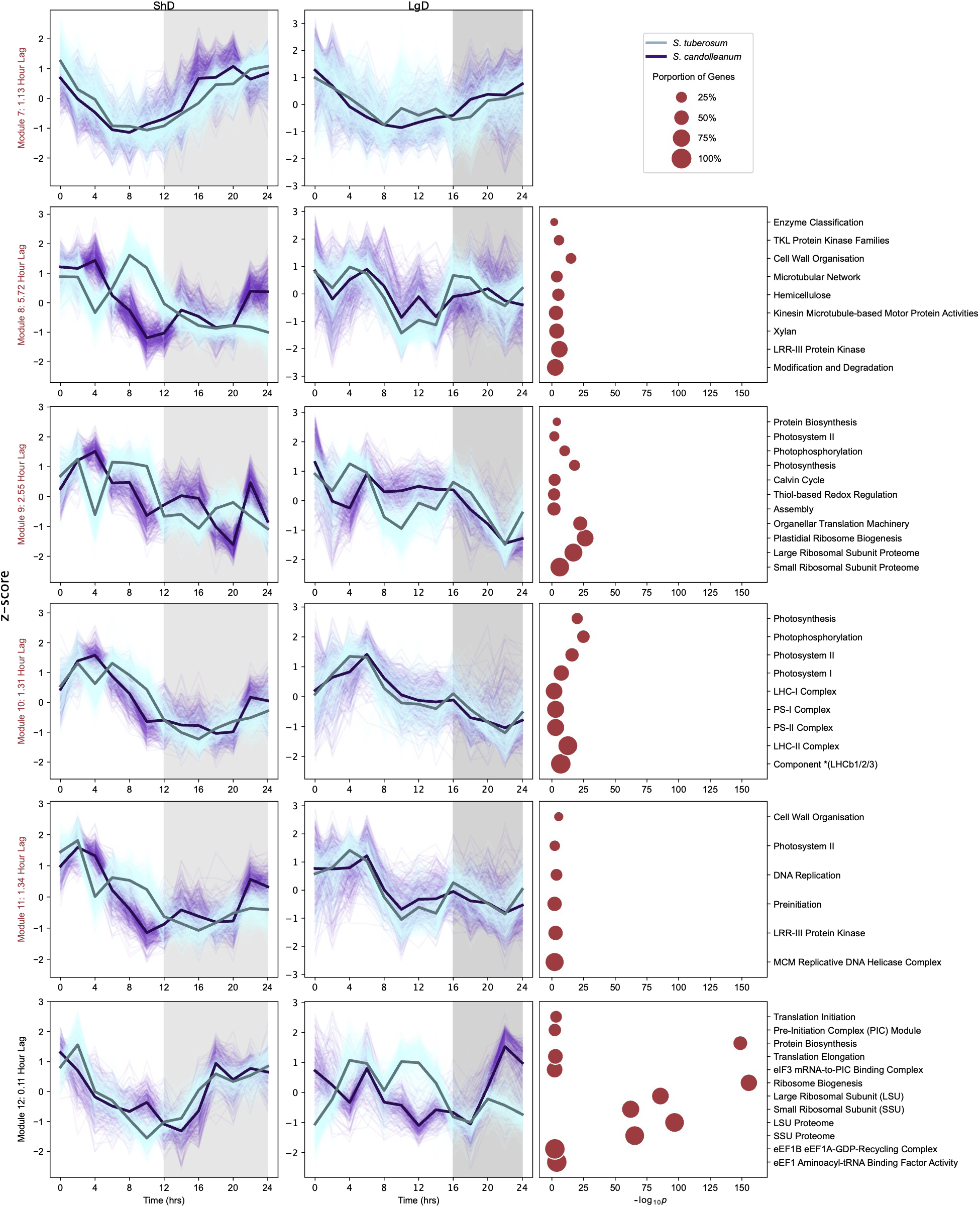
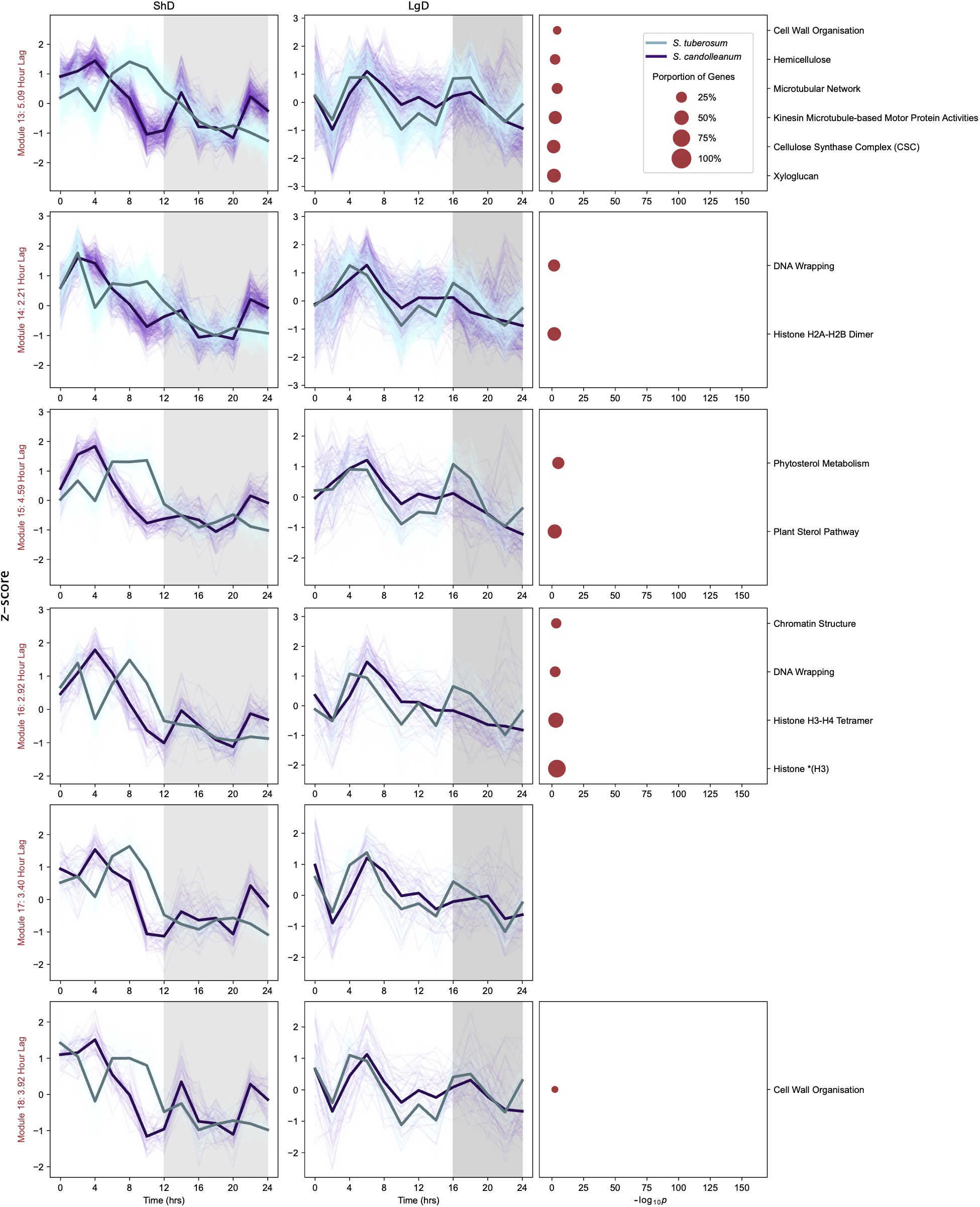
Expression patterns and mapman enrichment of genes within modules. Expression in *S. tuberosum* (blue) and *S. candolleanum* of genes within modules under short days (left) and long days (center). Both the individual expression patterns (thin, lighter lines) and mean expression patterns (thick, dark lines) are shown for both species. Right, Mapman enrichment for the genes within each module are shown. The size of the circle represents the percentage of the number of genes found in any module with that mapman term that were found in that specific module, and the positioning on the x-axis represents the –log(*p*-value) of the enrichment. Grey boxes represent nighttime conditions. Modules with red labels are those that exhibit phase delays in *S. tuberosum*.

**Figure S8.**
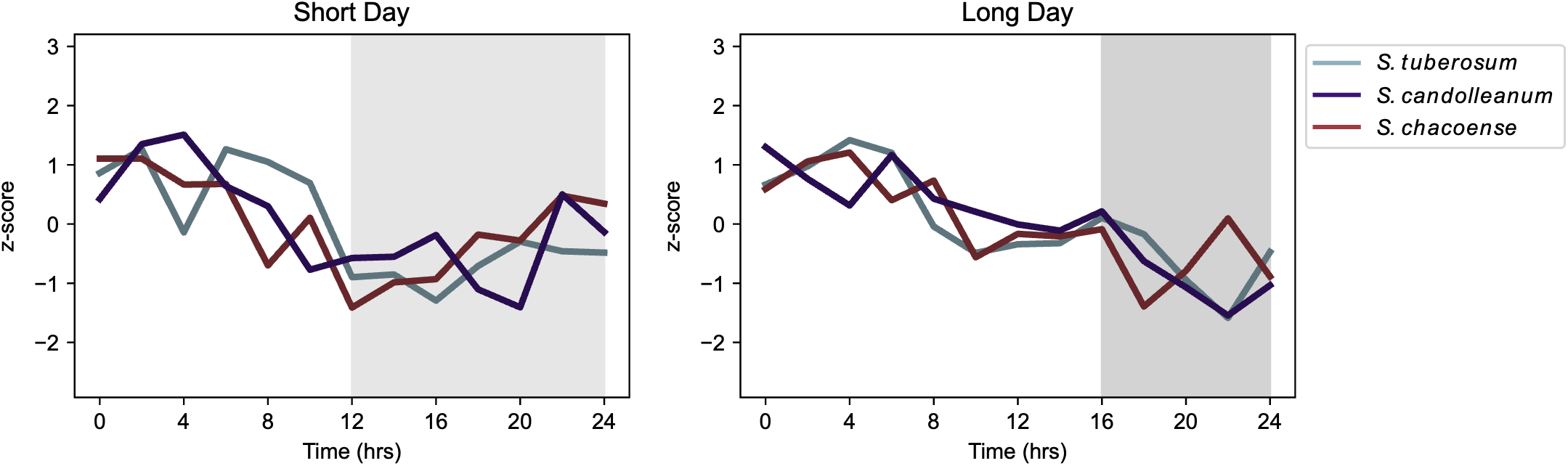
Photosynthesis genes in modules. Expression patterns of genes within modules 9, 10, and 11 that are annotated with the photosynthesis mapman term (term 17) under A. short day and B. long day conditions. Expression is shown for *S. tuberosum* (blue), *S. candolleanum* (purple) and *S. chacoense* (red). Both the individual expression patterns (thin, lighter lines) and mean expression patterns (thick, dark lines) are shown for all three species. Grey boxes represent nighttime conditions.

**Figure S9.**
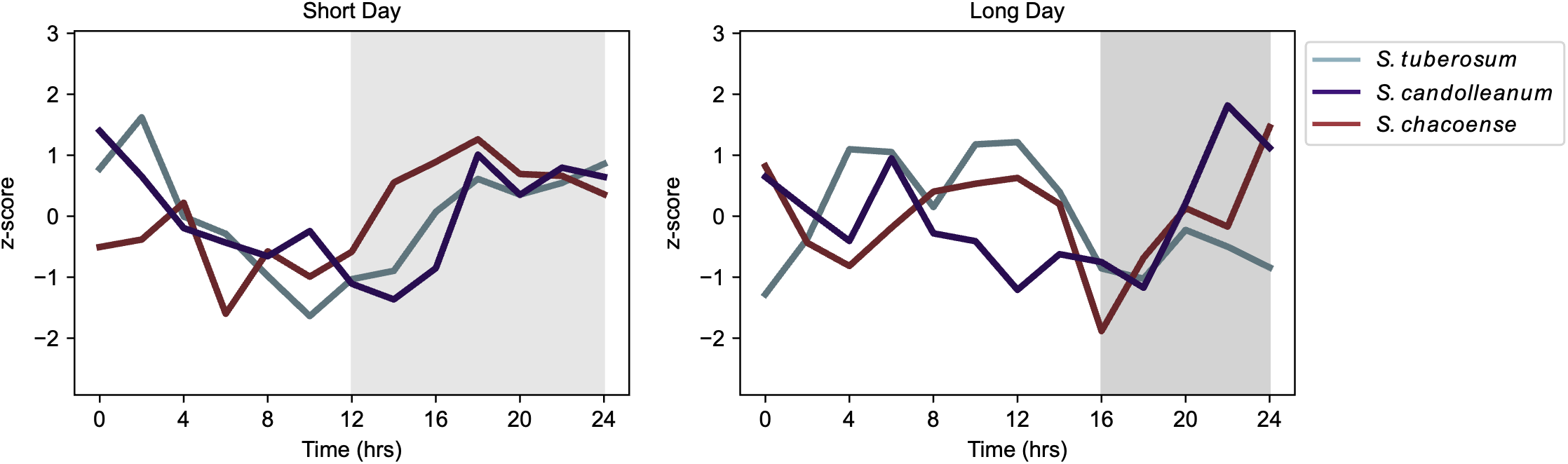
Protein biosynthesis genes in module 12. Expression patterns of genes within module 12 that are annotated with the protein biosynthesis Mapman term 17 under A. short day and B. long day conditions. Expression is shown for *S. tuberosum* (blue), *S. candolleanum* (purple) and *S. chacoense* (red). Both the individual expression patterns (thin, lighter lines) and mean expression patterns (thick, dark lines) are shown for all three species. Grey boxes represent nighttime conditions.

**Figure S10.**
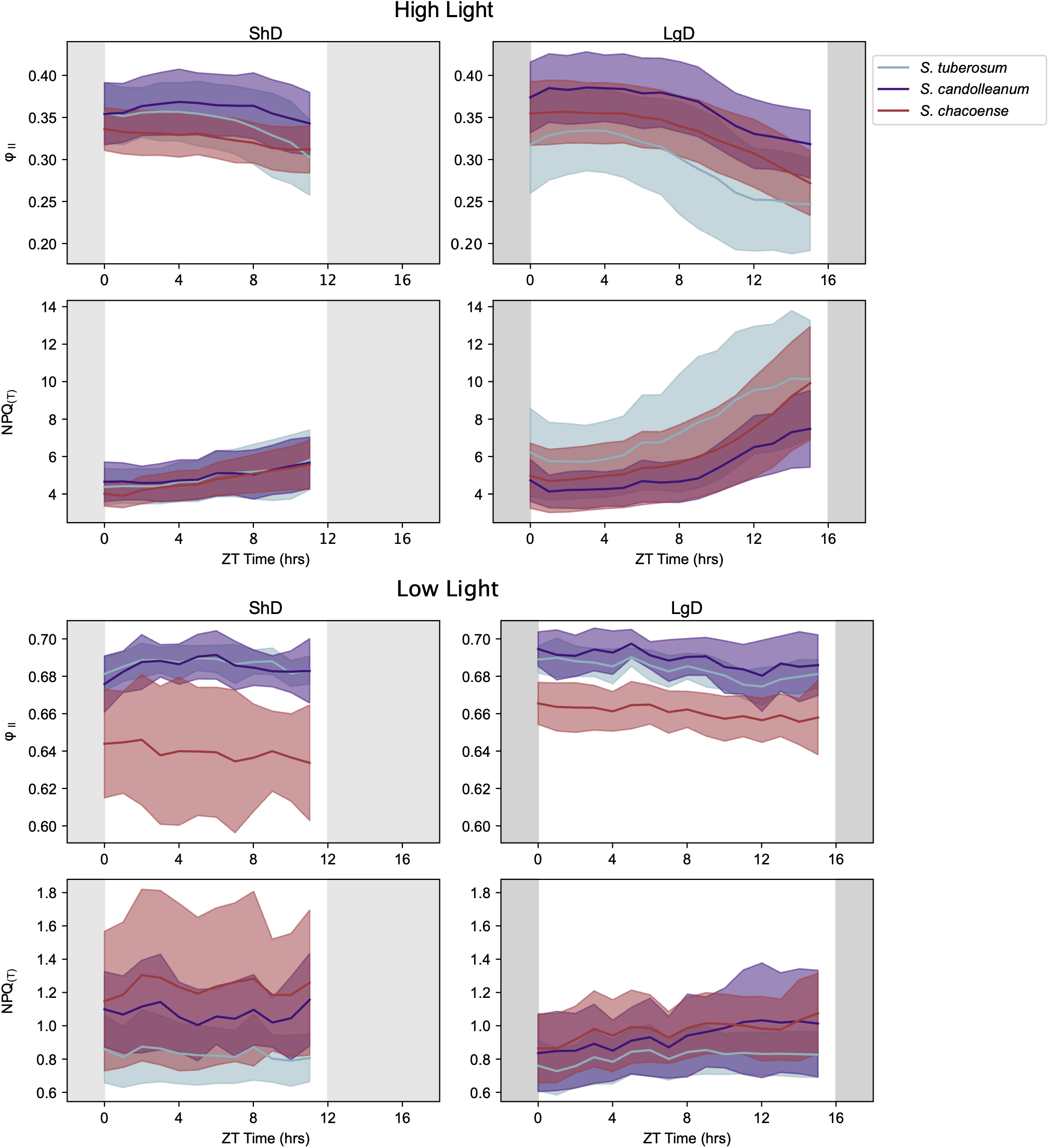
Differences in photosynthesis between species and photoperiods. Plots Φ₂ and NPQ(T) in 1000 mmol m^-2^ s^-1^ and 180 mmol m^-2^ s^-1^ light under short days and long days for *S. tuberosum* (blue), *S. candolleanum* (purple) *and S. chacoense* (red). Lines represent the mean of 16 independent samples while shaded regions represent the standard deviation. Grey boxes represent nighttime conditions.

**Figure S11.**
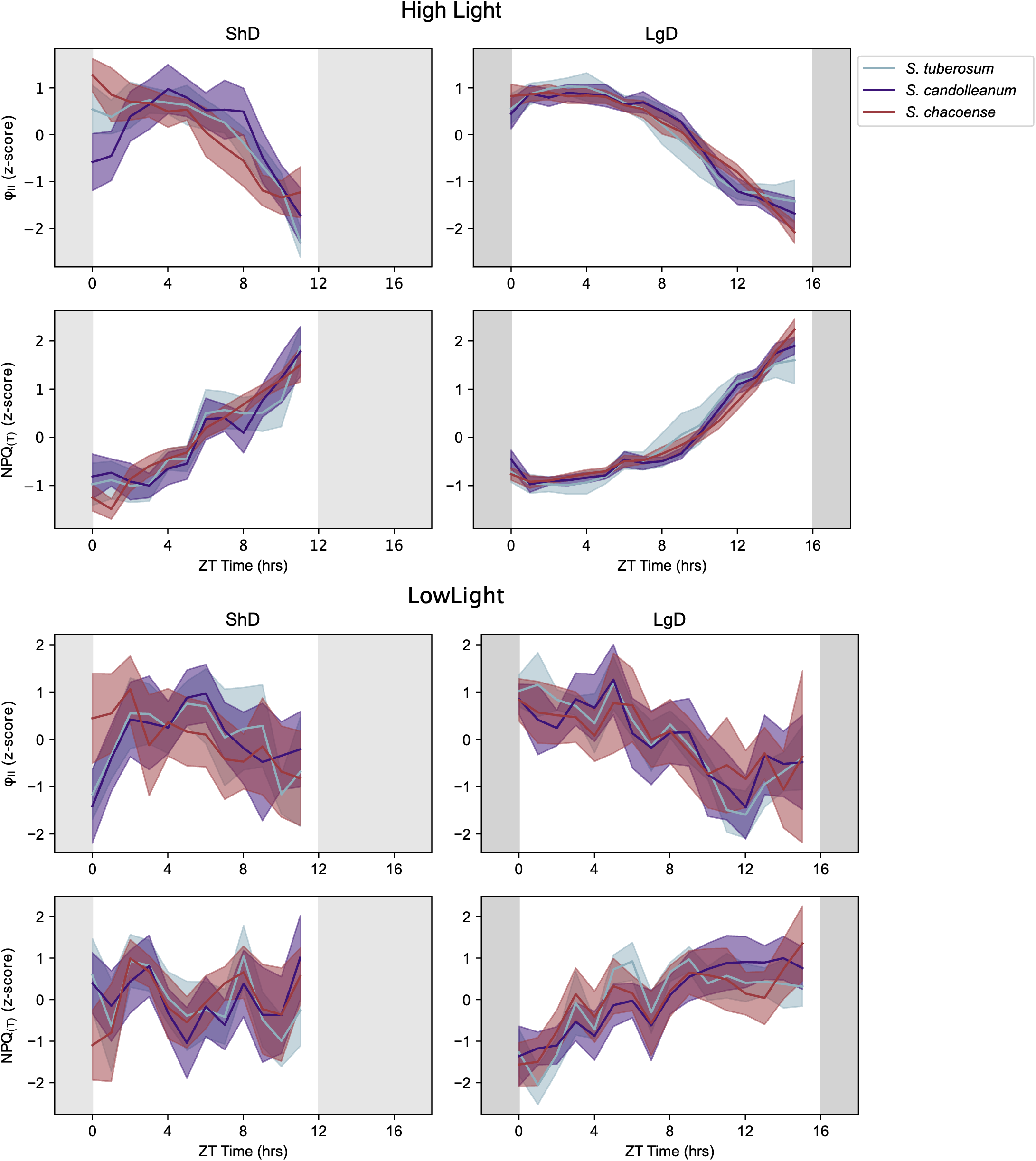
Differences in photosynthesis between species and photoperiods. Plots of the *z*-score of Φ₂ and NPQ(T) in 1000 mmol m^-2^ s^-1^ and 180 mmol m^-2^ s^-1^ light under short days and long days for *S. tuberosum* (blue), *S. candolleanum* (purple) *and S. chacoense* (red). Lines represent the mean of 16 independent samples while shaded regions represent the standard deviation. Grey boxes represent nighttime conditions.

**Figure S12.**
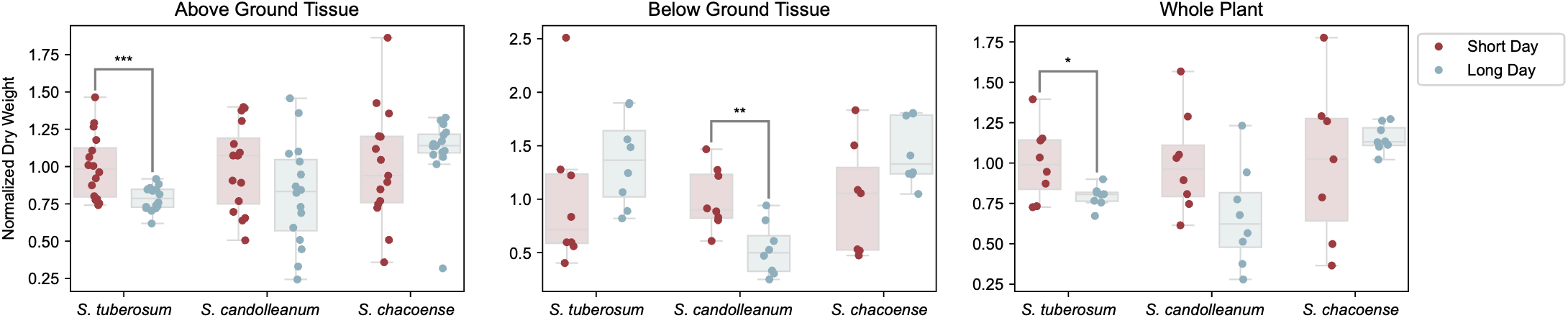
Differences in biomass between species and photoperiods. Plots of biomass for above ground tissue, below ground tissue, and whole plants under short days (red) and long days (blue) for all three species. Statistical significance was determined by a two-tailed Welch’s t-test. *: p<0.05, **: p<0.01, ***: p<0.001

**Figure S13.**
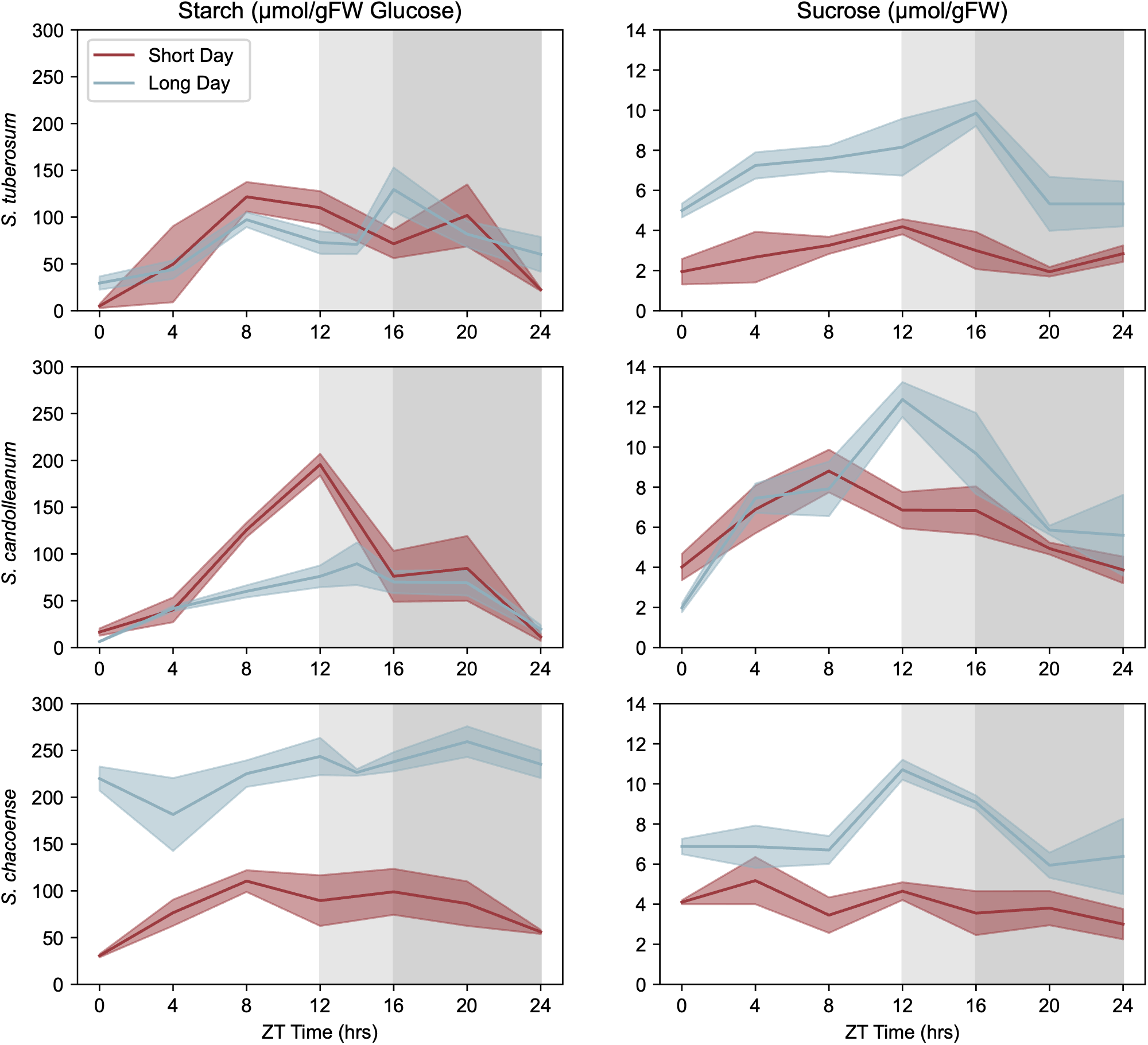
Differences in carbohydrate metabolites between species and photoperiods. Plots of starch and sucrose content for *S. tuberosum*, *S. candolleanum*, and *S. chaceoense* under short days (red) and long days (blue) for all three species. Lines represent the mean of six independent replicates while shaded regions represent the standard error. Grey boxes represent nighttime conditions under short days (light grey) and long days (dark grey).

**Figure S14.**
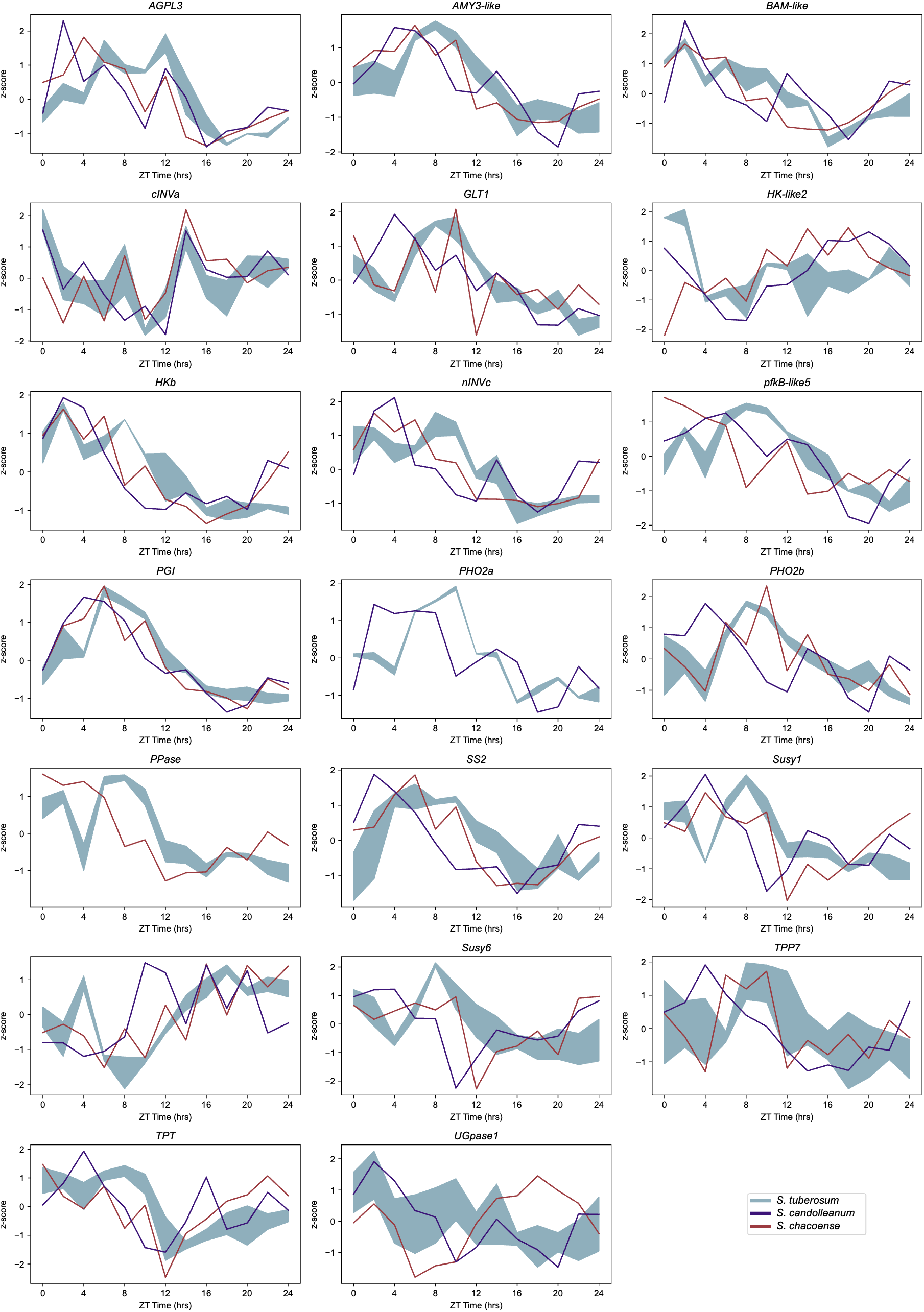
The expression patterns of selected carbohydrate genes in *S. tuberosum* (blue), *S. candolleanum* (purple), and *S. chacoense* (red) under short day conditions. *S. tuberosum* values are the mean ± standard deviation of all rhythmic alleles. Grey boxes represent nighttime conditions.

**Figure S15.**
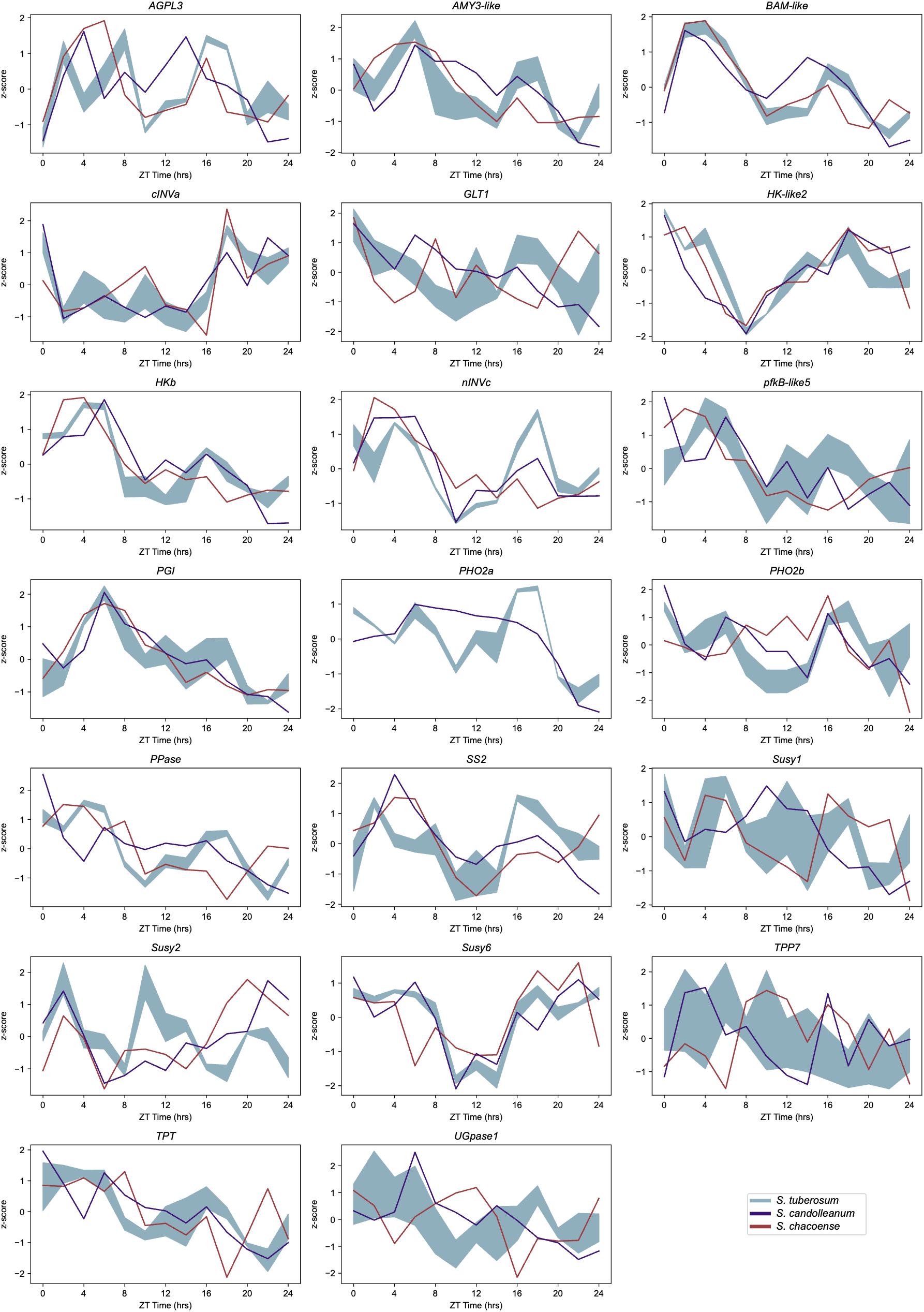
The expression patterns of selected carbohydrate genes in *S. tuberosum* (blue), *S. candolleanum* (purple), and *S. chacoense* (red) under long day conditions. *S. tuberosum* values are the mean ± standard deviation of all rhythmic alleles. Grey boxes represent nighttime conditions.

**Figure S16.**
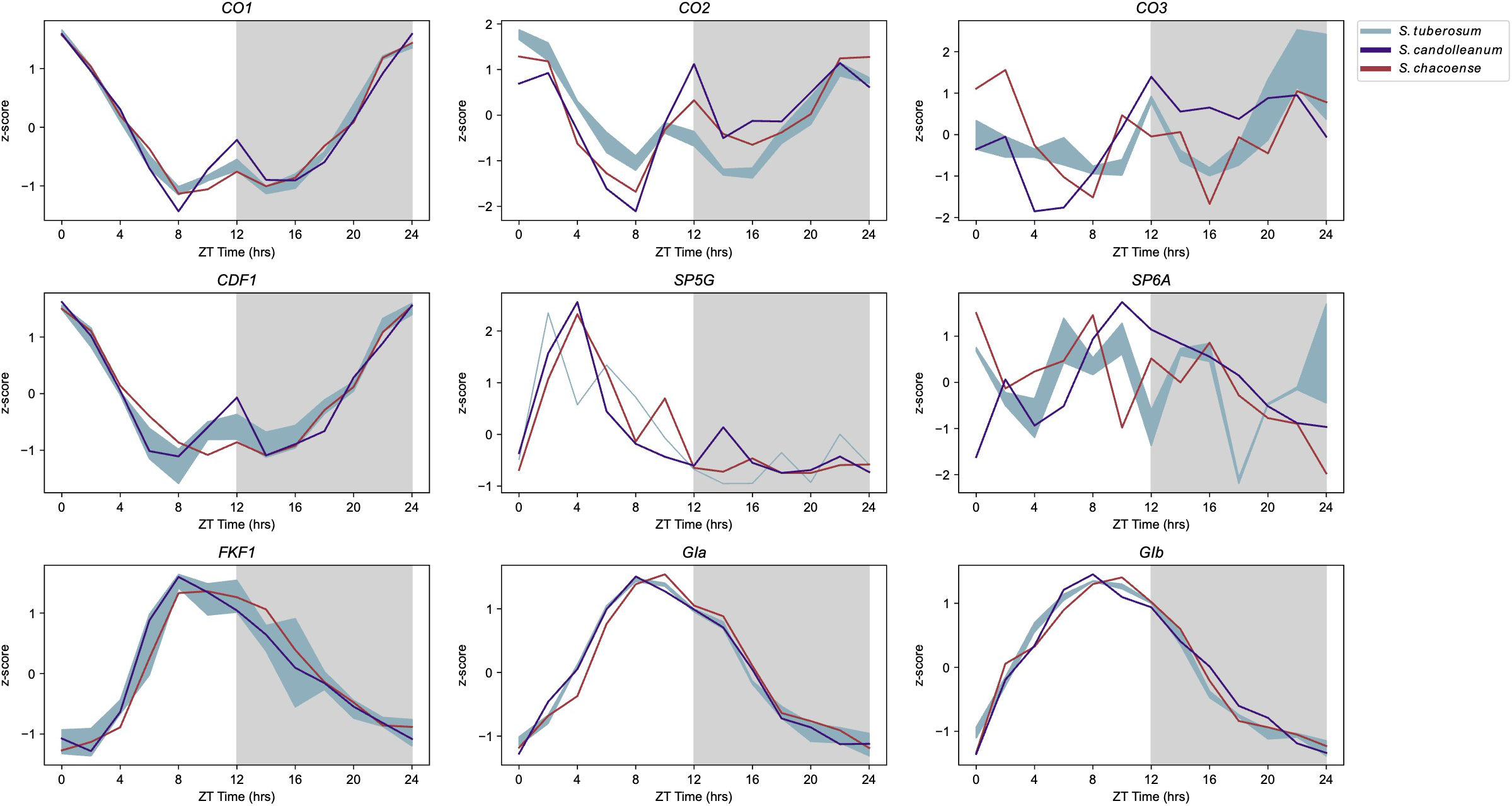
The expression patterns of photoperiod-related tuberization genes in *S. tuberosum* (blue), *S. candolleanum* (purple), and *S. chacoense* (red) under short day conditions. *S. tuberosum* values are the mean ± standard deviation of all expressed alleles. Grey boxes represent nighttime conditions.

**Figure S17.**
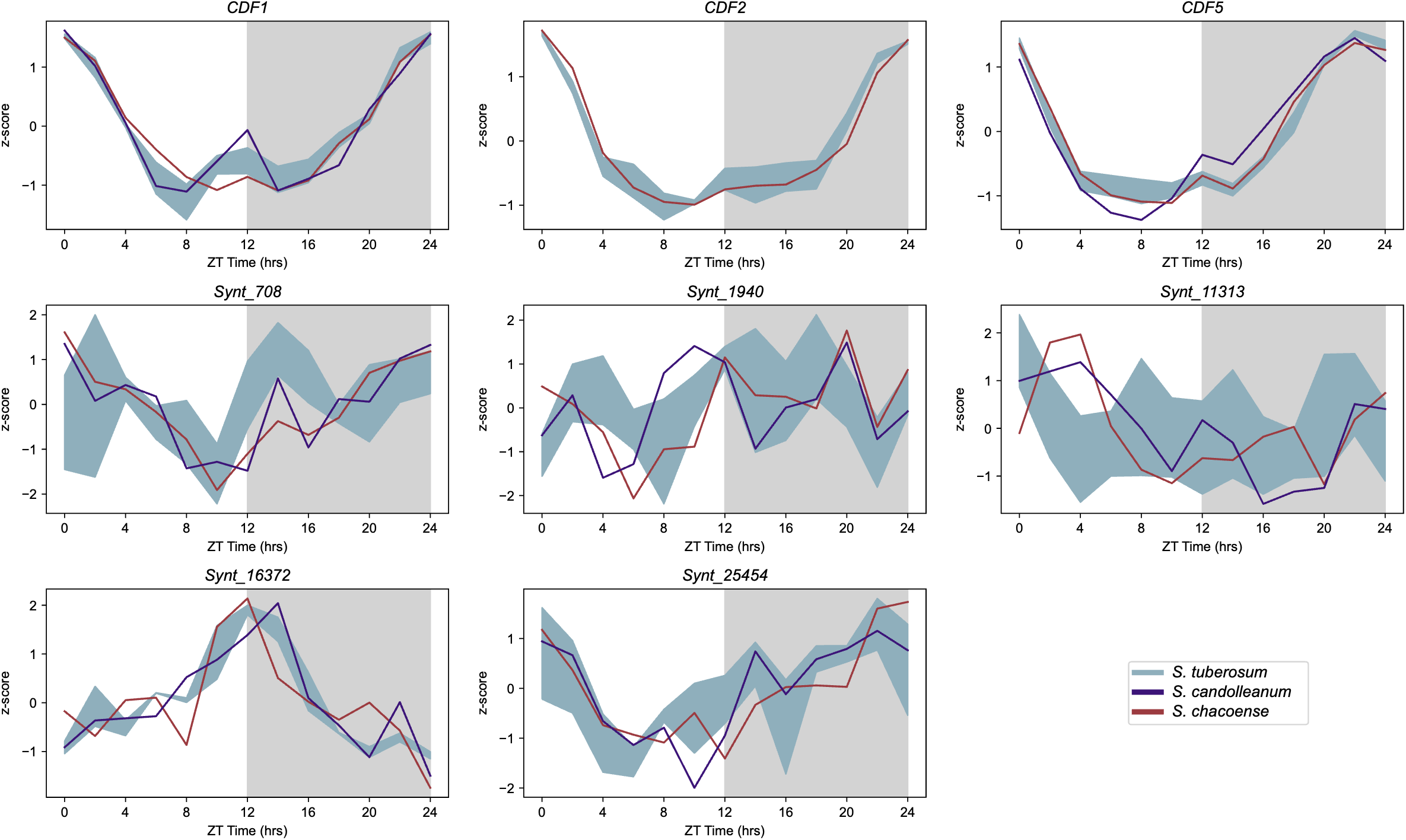
The expression patterns of TF group 4 genes in *S. tuberosum* (blue), *S. candolleanum* (purple), and *S. chacoense* (red) under short day conditions. *S. tuberosum* values are the mean ± standard deviation of all expressed alleles. Grey boxes represent nighttime conditions.

**Figure S18.**
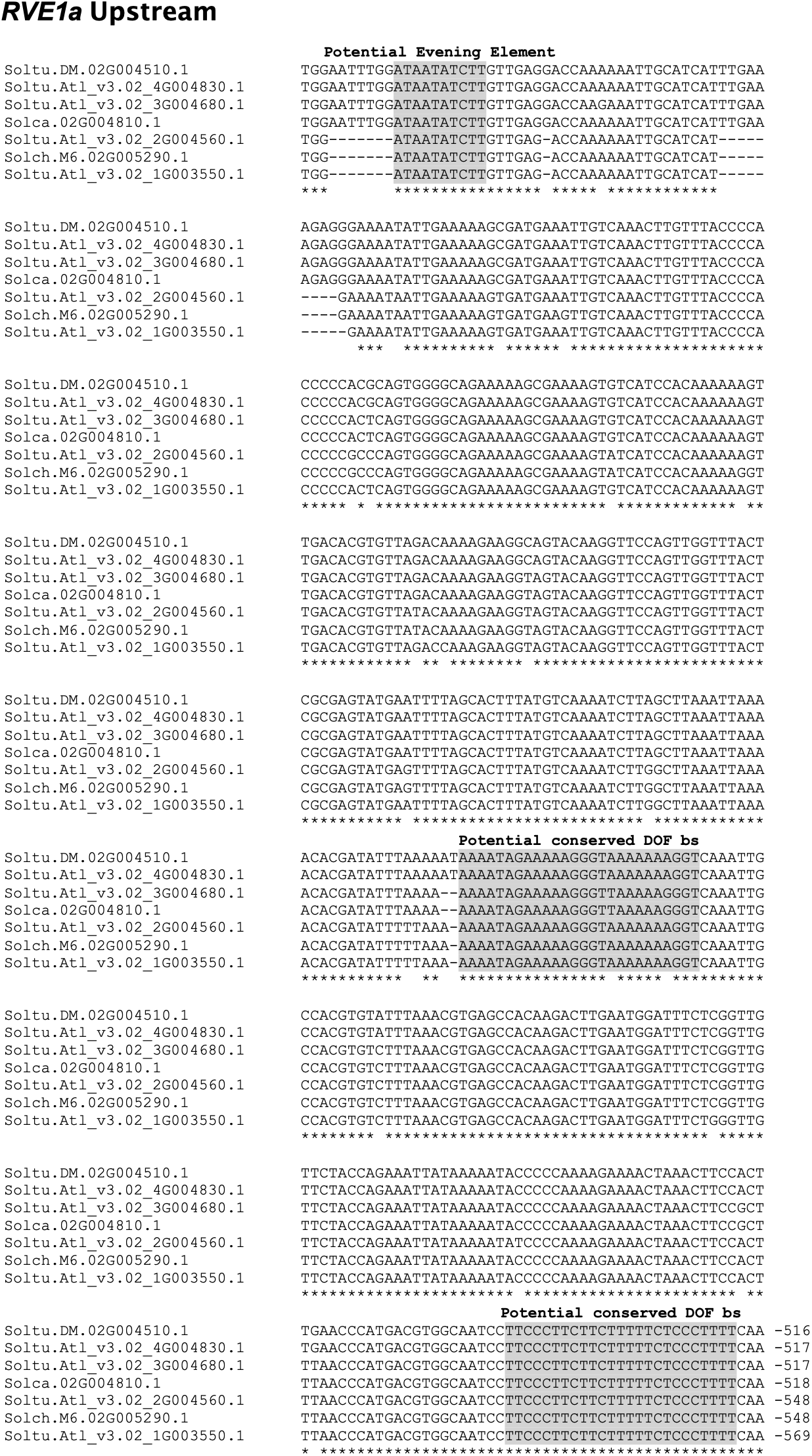
Alignment of RVE1a upstream sequences in potatoes. Potential cis-regulatory elements are highlighted in grey. Location is given with respect to annotated translational start side. Gene models are from *S. tuberosum* group Phureja DM1-3, *S. tuberosum* cv. Atlantic, *S. chacoense* and *S. candolleanum*. Sequences were aligned using ClustalW.

**Figure S19.**
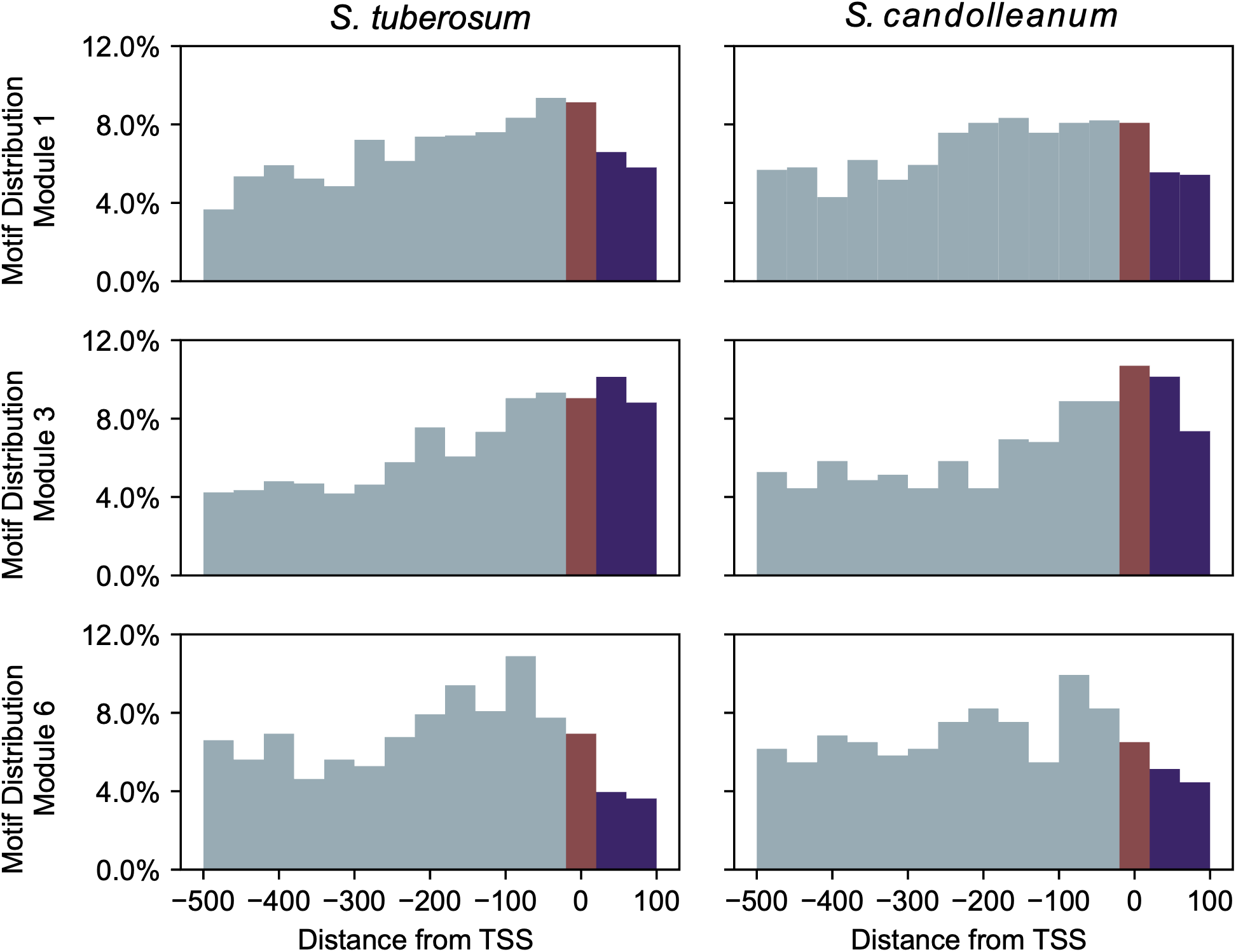
Histogram of the location of the Evening Element within the promoters of genes in module 1, 3 and 6 in *S. tuberosum* and *S. candolleanum.* Purple bars are those where the Evening Element is after the transcription start site, blue bars are those where the Evening Element is before the transcription start site, and red bars are those where the Evening Element contains the transcription start site.

## Supplemental tables

Table S1. Genome assembly data summary data.

Table S2. BUSCO analysis for *S. chacoense*.

Table S3. BUSCO analysis for *S. candolleanum*.

Table S4. Functional enrichment of genes rhythmic in all three potato species.

Table S5. Conserved circadian clock associated genes.

Table S6. Syntelogs in each S*. tuberosum*-*S. cadolleanum* coexpression modules.

Table S7. Carbohydrate genes in S*. tuberosum*-*S. cadolleanum* coexpression modules.

Table S8. Transcription factor genes in coexpression modules and their corresponding motifs.

Table S9. Gene regulatory network.

Table S10. Genes in coexpression module 11.

Table S11. Libraries used in assembly, annotation, and analysis of *Solanum chacoense* and *Solanum candolleanum*.

Table S12. Information on the RNA-seq data used in this study. Table S13. Primers used in this study.

## Supplemental large datasets

Dataset S1. List of syntelogs in *S. tuberosum* cv. Atlantic, *S. candolleanum*, *S. chacoense*, *S. tuberosum* group Phureja DM1-3.

Dataset S2. Short-day expression rlog for *S. tuberosum*. Dataset S3. Long-day expression rlog for *S. tuberosum*.

Dataset S4. Short-day expression rlog for *S. candolleanum*.

Dataset S5. Long-day expression rlog for *S. candolleanum*.

Dataset S6. Short-day expression rlog for *S. chacoense*.

Dataset S7. Long-day expression rlog for *S. chacoense*.

Dataset S8. Short-day tissue cycling parameters as determined per JTK for *S. tuberosum*.

Dataset S9. Long-day tissue cycling parameters as determined per JTK for *S. tuberosum*.

Dataset S10. Short-day tissue cycling parameters as determined per JTK for *S. candolleanum*.

Dataset S11. Long-day tissue cycling parameters as determined per JTK for *S. candolleanum*.

Dataset S12. Short-day tissue cycling parameters as determined per JTK for *S. chacoense*.]

Dataset S13. Long-day tissue cycling parameters as determined per JTK for *S. chacoense*.

Dataset S14. Functional annotation of *S. candolleanum* using MapMan.

Dataset S15. Functional annotation of *S. chacoense* using MapMan.

Dataset S16. Combined functional annotation for *S. tuberosum* cv. Atlantic, *S. candolleanum*, and *S. chacoense* syntelogs.

